# Investigating the coupled effects of stiffness and stretch on the trabecular meshwork cells using a hydrogel-integrated microfluidic system

**DOI:** 10.64898/2026.04.17.717863

**Authors:** Kanghoon Choi, Minju Kim, Monika Lakk, Fiona S. McDonnell, David Krizaj, Jungkyu Kim

**Affiliations:** Department of Mechanical Engineering, University of Utah, Salt Lake City, UT 84112; Department of Ophthalmology & Visual Sciences, University of Utah School of Medicine, Salt Lake City, UT 84112; Department of Biomedical Engineering, University of Utah School of Medicine, Salt Lake City, UT 84112; Department of Pharmacology and Toxicology, University of Utah School of Medicine, Salt Lake City, UT 84112

**Keywords:** Glaucoma, Trabecular Meshwork, Hydrogel-integrated Microfluidics, Organ-on-a-chip, Mechanotransduction, Substrate Stiffness

## Abstract

Glaucoma is characterized by progressive stiffening of the trabecular meshwork (TM), which elevates intraocular pressure and contributes to tissue dysfunction. Although substrate stiffness and mechanical stimulation both regulate TM homeostasis, their combined effects remain poorly understood. Here, a hydrogel-integrated microfluidic platform is presented that enables simultaneous control of substrate stiffness via tunable gelatin methacryloyl (GelMA) hydrogels and equi-biaxial quasi-static stretch via hydraulic actuation. Finite element analysis validates the applied strain field, and optimized crosslinking ensures structural stability. Primary normal TM (nTM) and glaucomatous TM (gTM) cells cultured under coupled conditions exhibit selective mechanotransduction dysregulation rather than global mechanosensory impairment. While nTM cells dynamically regulate α-smooth muscle actin (α-SMA), myocilin (MYOC), matrix metalloproteinase-2 (MMP2), and collagen type I (COL1), gTM cells display constitutively elevated α-SMA, loss of mechanical regulation of MMP2, and impaired stretch-mediated COL1 suppression, while retaining stiffness-dependent focal adhesion kinase and MYOC sensitivity. Key differences between normal and glaucomatous cells emerge only under combined stiff and stretched conditions, underscoring the importance of coupled mechanical cues in revealing disease-relevant phenotypes. These findings implicate tissue stiffening in selective pathway dysregulation and highlight mechanotransduction-targeted therapeutic strategies.

## 1. Introduction

Glaucoma affects over 76 million people worldwide, causing irreversible vision loss through progressive optic nerve degeneration. By 2040, this number is projected to exceed 111 million ^[1]^, with primary open-angle glaucoma (POAG) accounting for most cases ^[2]^. The disease originates from impaired aqueous humor outflow through the trabecular meshwork (TM), where pathological tissue stiffening is thought to create a progressive pathological cascade: increased stiffness elevates outflow resistance, raising intraocular pressure (IOP), which may further stiffen the tissue through mechanotransduction pathways ^[3]^. In healthy eyes, the TM maintains a compliant structure with stiffness of approximately 4 kPa, facilitating dynamic regulation of aqueous humor drainage. During glaucoma progression, tissue stiffness can increase up to 20-fold, exceeding 80 kPa in advanced disease ^[4]^. This pathological stiffening occurs heterogeneously, creating mechanical gradients that expose cells to complex, non-uniform stress fields during IOP fluctuations ^[5]^.

In addition to pathological stiffening, TM cells experience mostly quasi-static stretch from baseline IOP, with smaller fluctuations caused by ocular pulse and ciliary muscle activity ^[6]^. Early histological observations demonstrated that raising perfusion pressure from 8 to 30 mmHg elongates trabecular beam elements by up to ∼50%^[7]^, although the effective strain transmitted to individual TM cells is expected to be substantially lower due to the hierarchical architecture of the meshwork ^[8]^. Experimental studies have accordingly applied substrate elongation spanning small-amplitude cyclic stretch (∼0.45% at 1 Hz) to mimic ocular pulsations, up to larger magnitudes (∼20%) intended to probe TM mechanobiology and model pathological loading in vitro^[9]^. Such stretch activates mechanosensitive ion channels, integrin-mediated mechanotransduction, driving actin polymerization, focal adhesion remodeling with phosphorylation of focal adhesion kinase (pFAK), and downstream ECM secretion^[8a^, ^10]^. Characteristic TM responses include myofibroblast differentiation with elevated α-smooth muscle actin expression (α-SMA)^[11]^, upregulation of glaucoma-associated myocilin (MYOC)^[12]^, increased matrix metalloproteinase-2 (MMP2) activity^[13]^, altered collagen type I (COL1) organization^[14]^, and cytoskeletal reorganization with focal adhesion remodeling^[15]^. Consistent with this framework, 10% mechanical stretch of human TM cells rapidly upregulates MYOC mRNA within 8–24 h^[12]^, while cyclic stretch paradigms preferentially drive MMP-mediated matrix turnover and outflow-pathway remodeling^[8b^, ^16]^. More recent work further shows that these stretch responses in TM cells require parallel activation of TRPV4 channels, RhoA/ROCK, and integrin–FAK signaling, linking ion-channel mechanosensing directly to classical mechanotransduction pathways^[17]^.

Despite advances in understanding the mechanobiology of glaucoma, a fundamental gap persists in elucidating how TM cells integrate distinct mechanical cues, including substrate stiffness and quasi-static stretch, to regulate disease-associated responses. Current in vitro models typically examine these factors independently, failing to capture their synergistic effects in a physiological context. For example, static culture on stiff substrates induces myofibroblast transformation ^[11b, 18]^, while stretch promotes ECM remodeling ^[14b, 19]^; yet the combined effects of stiffness and stretch remain largely unexplored, despite significant progress in organ chips for mechanobiology studies. Existing organ chips face several critical limitations, including the inability to combine tunable hydrogel substrates with mechanical actuation, restriction to uniaxial or biaxial deformations that do not replicate physiological equi-biaxial stress states, and poor hydrogel integration under quasi-static loading^[20]^. Previous PDMS-based stretching platform achieved mechanical deformation but lacked tunable stiffness, whereas hydrogel-based platforms provided substrate control without enabling mechanical stimulation ^[21]^.

To address these critical limitations, we present a novel hydrogel-integrated microfluidic platform enabling simultaneous control of substrate stiffness and equi-biaxial mechanical stretch. Our approach integrates tunable gelatin methacryloyl (GelMA) hydrogel substrates spanning the pathophysiological stiffness range with hydraulic actuation for precise mechanical stretch. The system addresses key technical challenges through advanced surface chemistry for robust hydrogel integration and a multi-layer architecture design that maintains mechanical integrity under static loading. Using this platform, we investigate how substrate stiffness creates a mechanically permissive environment that amplifies cellular responses to stretch, potentially contributing to a pathological feedback loop associated with glaucoma progression. Our findings reveal previously unrecognized mechanobiological mechanisms whereby coupled mechanical stimuli synergistically regulate TM cell behavior associated with pathology. Notably, we demonstrate that appropriate mechanical modulation can partially modulate disease-associated markers in glaucomatous cells, suggesting avenues for further investigation of therapeutic approaches.

## 2. Results

### 2.1 Hydrogel-Integrated Microfluidic System

We designed a hydrogel-integrated microfluidic chip enabling multiple experiments under controlled stiffness and mechanical stretch (**Fig. 1A**). The chip comprises a glass substrate, five PDMS layers, and a GelMA layer. A cross-shaped microfluidic channel enables the simultaneous incorporation of four homogeneous GelMA hydrogels within 4 mm-diameter wells. These are nested within 8.5 mm wells, which are designed to hold 8 mm circular coverslips for preparing slab gels. A hydraulic pressure channel enables concurrent equi-biaxial inflation of the four GelMA/PDMS composite membranes (**Fig. 1B**). By integrating four identical units on a single chip, the platform enables simultaneous testing of four hydrogel stiffnesses across four stretch levels and, in principle, supports multiple culture configurations, including cells cultured on the hydrogel surface, embedded within the matrix, or positioned at the bottom of the wells. For functional validation, dye loading (**Fig. 1A**) demonstrated that all four chambers maintained independently controlled environments without cross-contamination. The PDMS–glass bond withstood repeated hydraulic actuation without leakage or delamination, and the membrane remained intact across the full operating range.

**Fig. 1.**
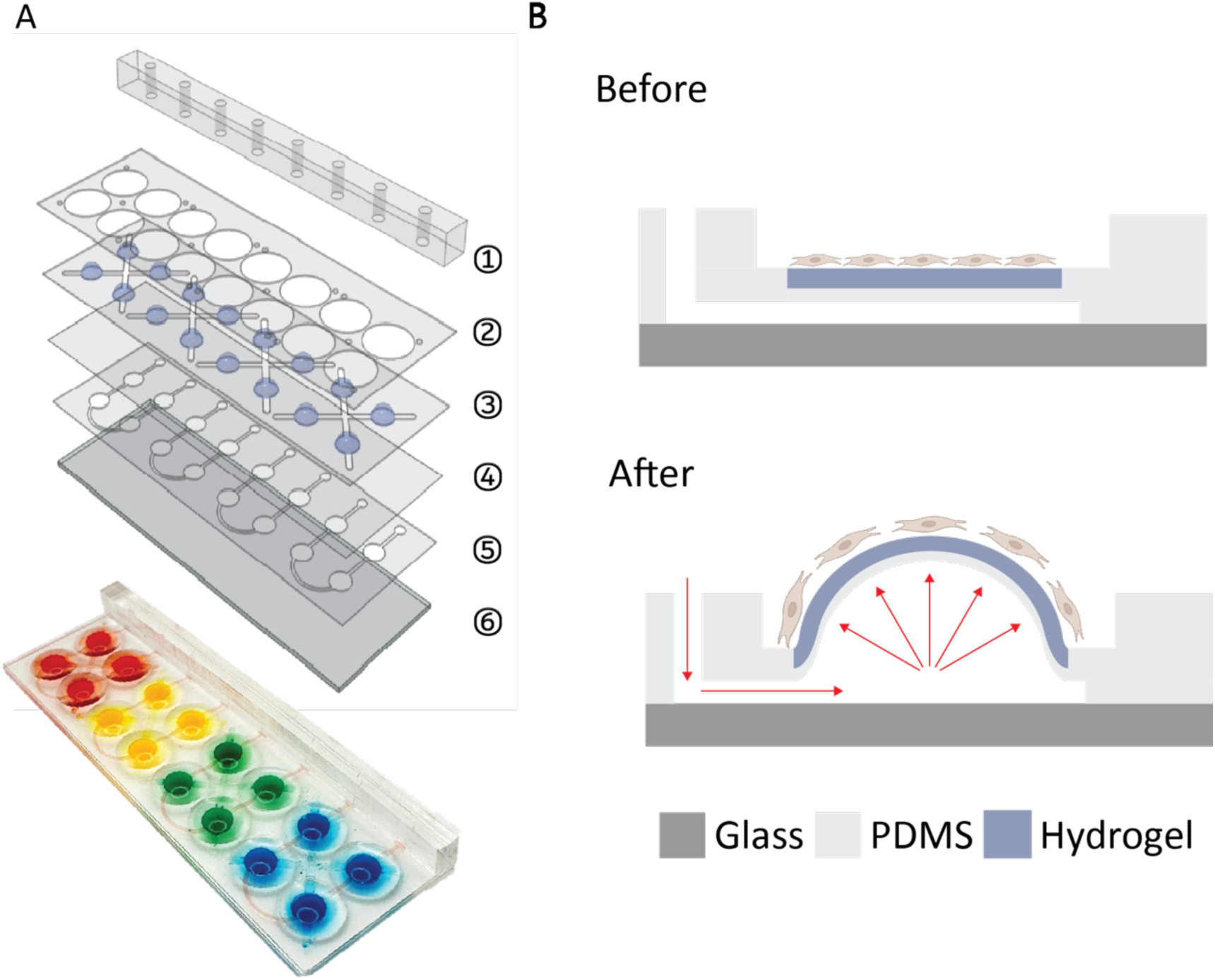
Hydrogel-integrated microfluidic system concept and design. (A) Schematic illustration and a photograph of the hydrogel-integrated microfluidic system showing the multi-layer architecture. (B) Cross-sectional view demonstrating hydraulic pressure application through the microfluidic channel to induce equi-biaxial stretch on the integrated hydrogel.

### 2.2 GelMA Hydrogel Stability and Mechanical Properties

The effects of UV photo-crosslinking duration and polymer concentration on GelMA hydrogel stability were systematically evaluated. **Fig. S1** shows that surface area analysis of 5% GelMA after 24 h in PBS revealed an inverse relationship between UV exposure time and shrinkage, with 60s producing 8.0 ±0.2% shrinkage and 150 s reducing it to 1.5 ±0.1%. Shrinkage plateaued beyond 150 s, with no significant changes at longer exposures, establishing 150 s as the optimal crosslinking time. **Fig. S2** shows that Young’s modulus increased with polymer concentration, measuring 1.23 ±0.21 kPa for 5%, 12.25 ±1.52 kPa for 10%, and 21.47 ±1.42 kPa for 20%.

### 2.3 Mechanical Characterization and Numerical Validation

To control strain via the hydraulic channel, we first determined the volume–strain relationship for the PDMS membrane and the PDMS/GelMA integrated layers. **Fig. 3A** shows the inflation of PDMS alone and the PDMS/GelMA composite layers, with the latter deforming conformally without any detachment. Strain calibration revealed excellent linearity between applied volume and linear strain for all layers (R² =0.99, **Fig. 3B**). The PDMS and PDMS/5% GelMA layers achieved up to ∼50% linear strain, whereas the PDMS/20% GelMA layer reached ∼40%, reflecting the higher crosslink density of the stiffer hydrogel. To evaluate the maximum operating volume, we performed burst testing and found that most layers could withstand up to 90 µL without leakage or delamination (**Fig. 3C**).

**Fig. 2.**
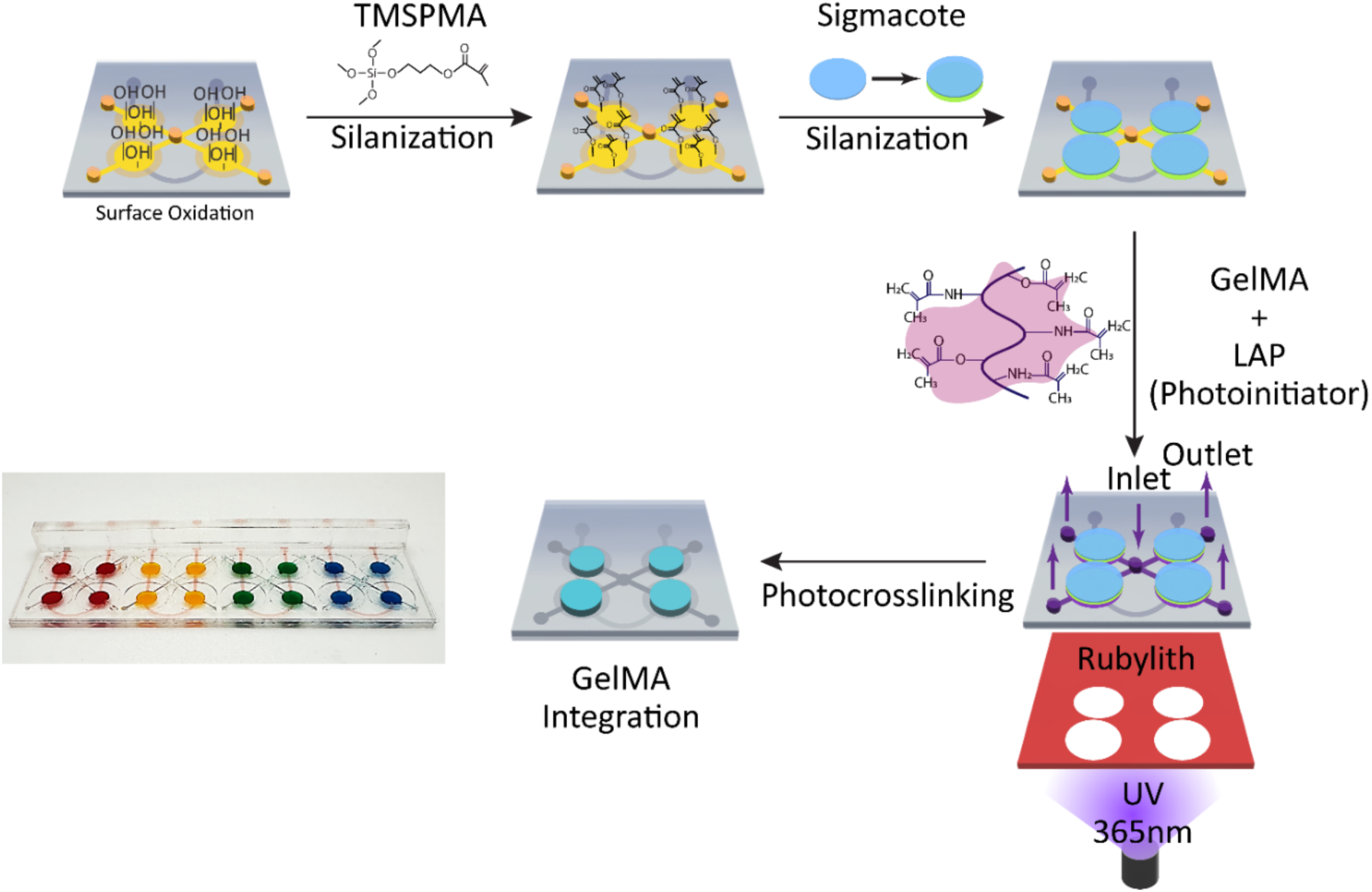
Surface modification and hydrogel integration. The PDMS surface was treated with oxygen plasma to form a hydroxyl group (OH-PDMS). 10% TMSPMA was treated to form a methacrylate functional group on the surface (TMSPMA-PDMS). A Sigmacote-coated coverslip was placed on the TMSPMA-PDMS to form an instant microfluidic channel. A patterned Rubylith film was placed on the bottom of the microfluidic system. GelMA hydrogel was introduced from the center inlet to form four hydrogels simultaneously. UV was exposed from the bottom of the chip. Coverslips were removed to create a flat hydrogel surface.

**Fig. 3.**
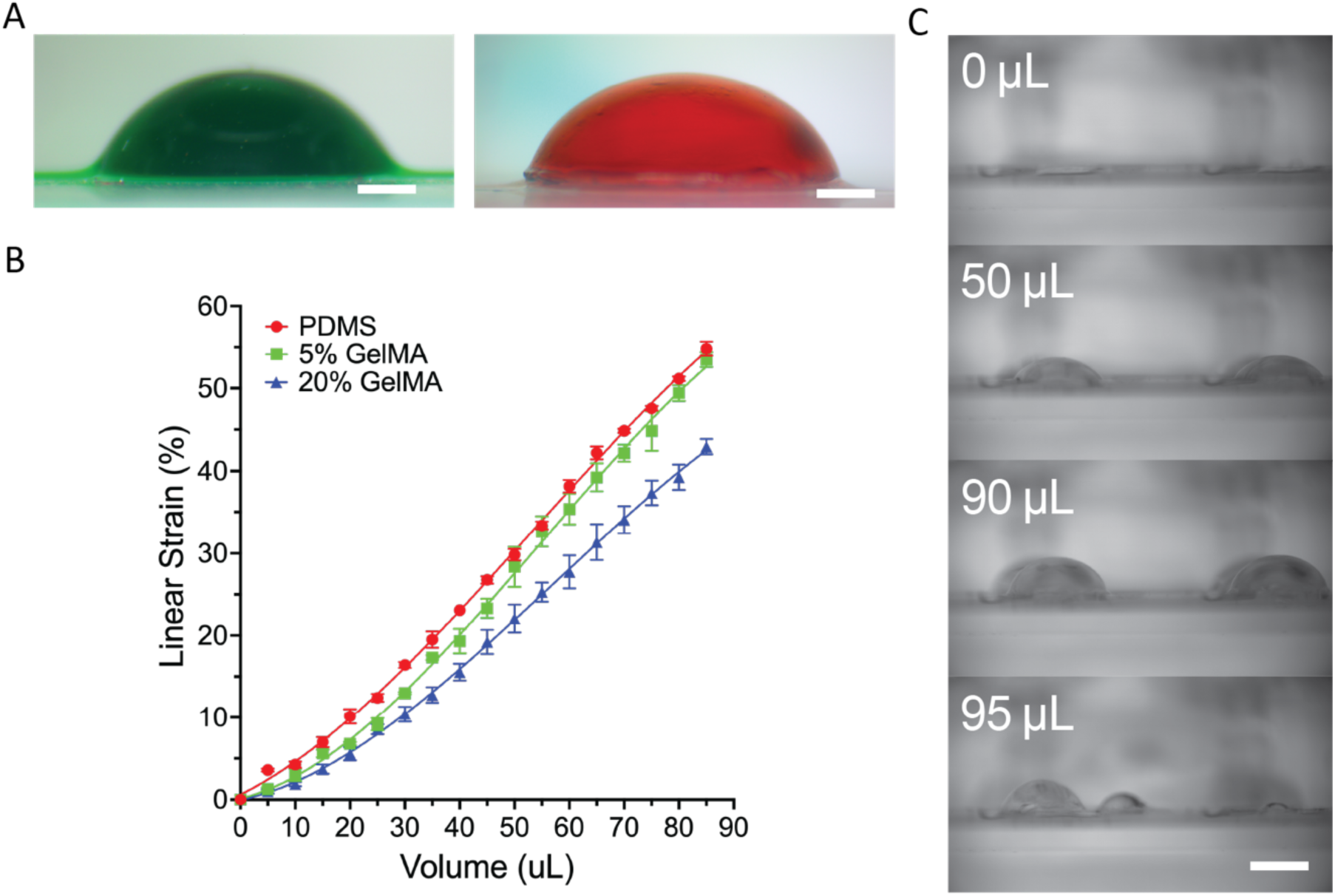
Mechanical characterization of PDMS and GelMA hydrogels for the strain analysis and bursting strength test. (A) Representative photographs of a PDMS substrate (left, green) and a GelMA hydrogel (right, red) within the experimental system. (B) Linear strain percentage as a function of applied fluid volume (µL) for PDMS, 5% GelMA, and 20% GelMA substrates. Data points represent mean ± standard deviation (n=3 for each material). (C) Sequential images illustrate the bursting strength test, showing increasing membrane deformation at 0 µL, 50 µL, 90 µL, and 95 µL of applied fluid volume. The membrane ruptured at volumes exceeding 95 µL.

Based on the hydrogel characterizations, we developed an FEA model to predict the mechanical behavior of the PDMS and PDMS/GelMA layers using Mooney–Rivlin and Veronda–Westmann constitutive models. Simulated peak dome deflections closely matched experimental measurements for all layers: PDMS (1.738 vs. 1.731 mm), PDMS/5% GelMA (1.927 vs. 1.927 mm), and PDMS/20% GelMA (1.704 vs. 1.703 mm), with the corresponding material constants summarized in **Table S1** (PDMS: C₁ =38.5 kPa, C₂ =2.93 kPa; 5% GelMA: C₁ =50 kPa, C₂ =0.42; 20% GelMA: C₁ =60 kPa, C₂ =0.46) ^[22]^. As expected, the PDMS/5% GelMA layer exhibited 13.1% greater deflection than the PDMS/20% GelMA layer, reflecting the lower stiffness of the softer hydrogel (**Fig. S3A**).

Using these models, unique orthogonal hoop and meridional stresses were captured under 10% equi-biaxial stretch. **Fig. 4C** shows that hoop and meridional stress profiles along the dome surface revealed a spatially heterogeneous mechanical environment. At the dome apex (Top), hoop and meridional stresses were nearly equal across all formulations, producing an equi-biaxial stress state. Toward the mid-slope (Mid), the hoop-to-meridional stress ratio progressively diverged as hoop stress increased and meridional stress decreased, reaching maximum stress anisotropy at the edge (Edge). Effective stress profiles summarized these directional variations as a single scalar gradient (**Fig. 4D**), with PDMS exhibiting the steepest gradient (57.9 to 14.9 kPa), followed by PDMS/20% GelMA (22.6 to 5.0 kPa) and PDMS/5% GelMA (15.8 to 3.8 kPa). Integration of GelMA attenuated effective surface stress by approximately 3-4 folds relative to pure PDMS while preserving the spatial gradient pattern. Cross-sectional views of displacement, effective stress, and effective solid stress for each case shown in **Fig. S3** confirmed these trends, with the effective solid stress isolating the stress gradient within the hydrogel layer to visualize the mechanical environment directly experienced by cells on the surface.

**Fig. 4.**
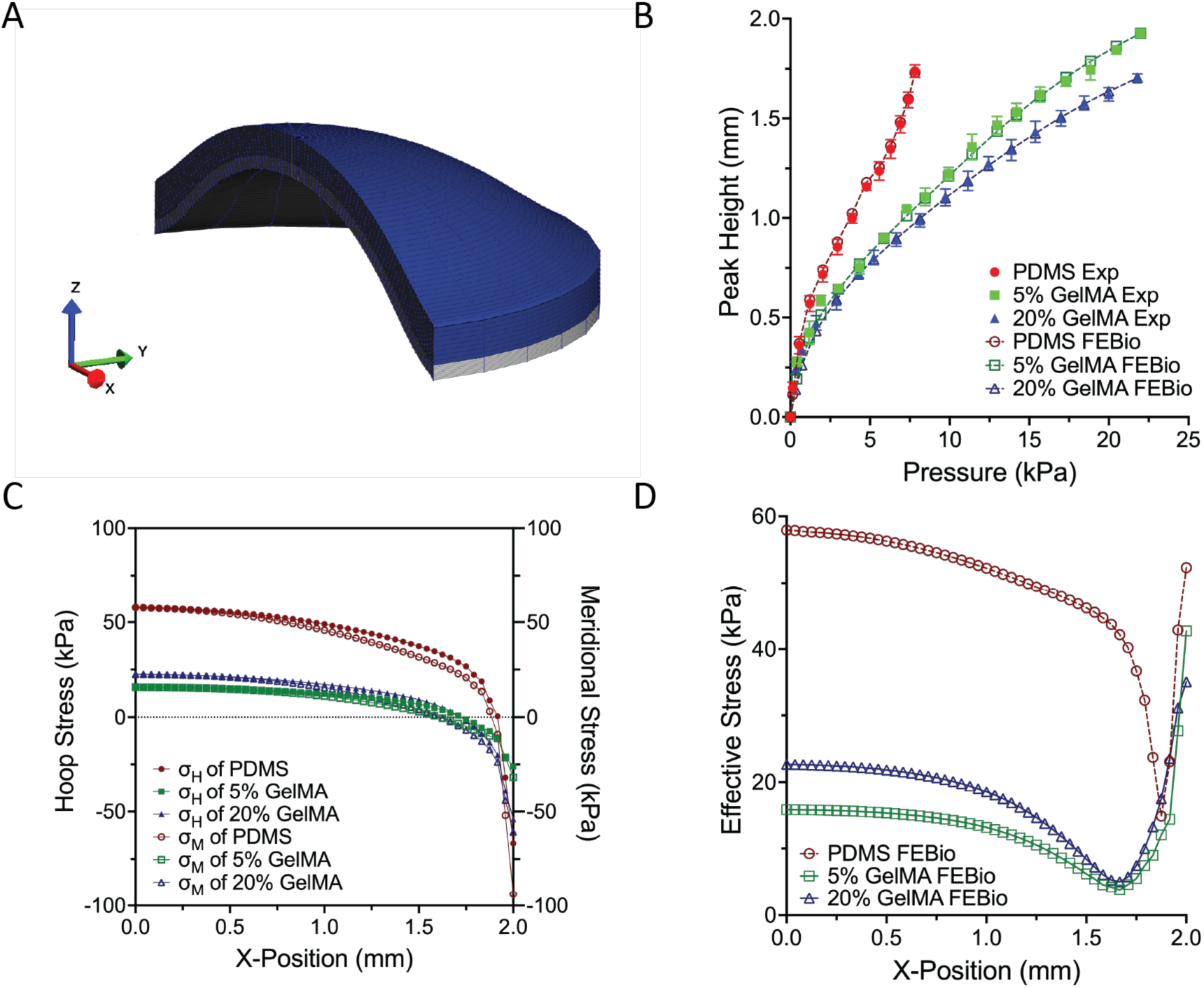
Finite element simulation and experimental validation of dome deformation and mechanical responses of GelMA and PDMS hemispherical constructs under pressure loading. (A) 3D finite element mesh model of a PDMS/20% GelMA hemispherical composite showing x-direction displacement under applied pressure. (B) Peak height as a function of applied pressure for PDMS, 5% GelMA, and 20% GelMA domes, comparing experimental measurements (solid markers) with finite element simulations (dashed lines). (C) Hoop and meridional stress distributions along the x-axis for each material system, illustrating the increase in stress magnitude with material stiffness. (D) Effective stress distribution along the x-axis, highlighting material-dependent stress localization; PDMS domes exhibit substantially greater deformation and higher peak stress compared to GelMA formulations.

### 2.4 Substrate Stiffness and Mechanical Stretch Regulate TM Cell Morphology

To investigate how mechanical stress influences normal TM(nTM) and glaucomatous TM (gTM) cell morphology, we quantified cell shape, spreading, and orientation under 10% equi-biaxial stretch across the three FEA-defined stress zones. **Fig. 5A** shows representative fluorescence images of nTM and gTM cells on 5% and 20% GelMA at the Top, Mid, and Edge regions. nTM cells progressively elongated from Top to Edge, reflecting increasing stress anisotropy, whereas gTM cells displayed region- and stiffness-dependent elongation patterns. **Fig. 5B** quantifies cell aspect ratio (minor axis/major axis; lower values indicate greater elongation). nTM on 5% GelMA showed significant decreases from Top to Mid (p ≤0.001) and Top to Edge (p ≤0.001), while cells on 20% GelMA followed a similar trend with reduced significance (Top vs. Mid: p ≤ 0.01; Top vs. Edge: p ≤ 0.05), suggesting partial constraint of elongation on stiffer substrates. gTM cells on 20% GelMA retained significant regional elongation (Top vs. Mid: p ≤0.05; Top vs. Edge: p ≤ 0.01), whereas gTM cells on 5% GelMA showed no significant differences, indicating impaired mechanosensing on compliant substrates. Cross-cell comparison at the Top region on 5% GelMA revealed that gTM cells were significantly more elongated than nTM cells (p ≤0.001). **Fig. 5C** presents cell spreading area. nTM cells increased spreading from Top to Edge on both 5% (p ≤0.01) and 20% GelMA (p ≤0.05), consistent with cytoskeletal remodeling in response to anisotropic stress. In contrast, gTM cells on 5% GelMA decreased spreading from Top to Edge (p ≤0.05), indicating divergent mechanoadaptive behavior. Cross-cell comparisons showed that gTM cells were larger than nTM cells at the Top region on both substrates (5% GelMA: p ≤0.01; 20% GelMA: p ≤0.001) and at the Mid region on 20% GelMA (p ≤0.01), but this trend reversed at the Edge on 5% GelMA, where nTM cells were larger (p ≤0.05). Cell orientation analysis (**Fig. 5A, S5**) confirmed region-dependent alignment. At the Top, cells were randomly oriented; at Mid, they aligned preferentially along the hoop stress direction; and at Edge, cells oriented along the meridional stress direction toward the dome center, consistent with local stress-driven cytoskeletal remodeling.

**Fig. 5.**
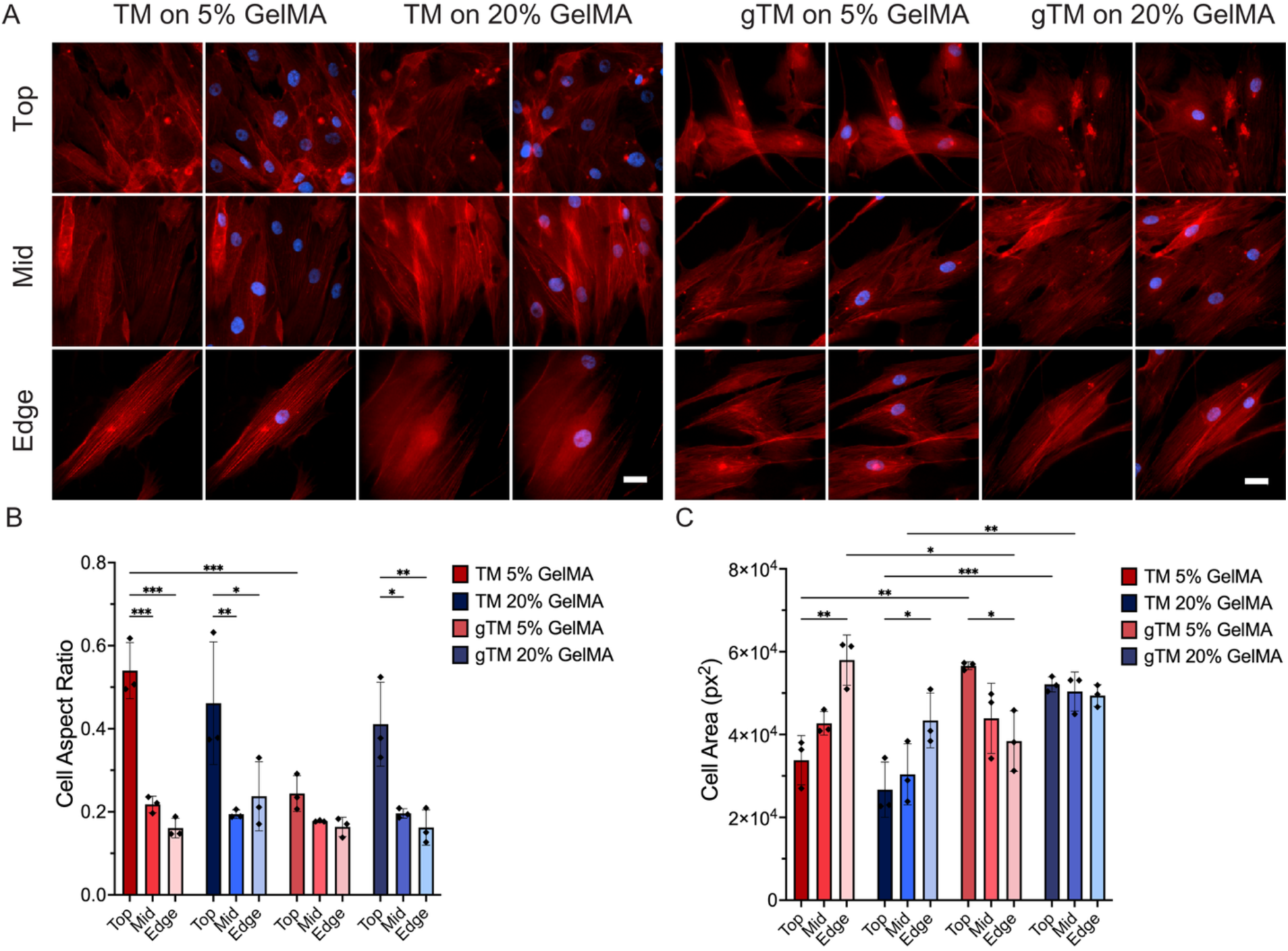
Morphological Characterization of nTM and gTM Cells on GelMA Hydrogels under Mechanical Stretch. (A) Representative immunofluorescence images of nTM and gTM cells on 5% and 20% GelMA hydrogels after 48 hours of culture, stained for F-actin (red) and nuclei (Hoechst, blue) across Top, Mid, and Edge dome regions. (B) Cell aspect ratio and (C) cell spreading area, quantified by dome region under 10% stretch. nTM cells exhibited greater aspect ratio changes across regions, indicating higher mechanosensitivity compared to gTM cells. nTM cells on both substrates and gTM cells on stiff substrates elongated progressively from Top to Edge regions, whereas gTM cells on compliant substrates lost significant regional elongation differences; nTM cells showed increased spreading area while gTM cells on compliant substrates displayed decreased spreading area, suggesting divergent mechanoadaptive responses. Data are presented as mean ± standard deviation (*p < 0.05, **p < 0.01, ***p < 0.001). Scale bar: 50 µm.

### 2.5 Phenotypic Markers of Glaucomatous Transformation

To investigate the impact of mechanical stress on phenotype, we examined α-SMA and MYOC expression in both nTM and gTM cells cultured on 5% and 20% GelMA hydrogels under no stretch or 10% equi-biaxial stretch. **Fig. 6A** (left panels) shows representative immunofluorescence images of α-SMA expression. In nTM cells, α-SMA signal was minimal on compliant substrates under static conditions, with sparse and weakly organized stress fibers. Increasing stiffness to 20% GelMA produced more prominent α-SMA-positive stress fibers, and the combination of 20% GelMA with 10% stretch yielded the most intense signal, with well-defined α-SMA incorporation into thick stress fibers that co-localized strongly with F-actin. In contrast, gTM cells exhibited robust α-SMA-positive stress fibers across all conditions, including on compliant substrates without stretch, visually confirming a constitutively activated myofibroblast phenotype. **Fig. 6B-a** quantifies these observations. In nTM cells, α-SMA was significantly modulated by both substrate stiffness and stretch. Increasing stiffness from 5% to 20% GelMA elevated α-SMA under no stretch (p ≤ 0.001) and 10% stretch (p ≤ 0.01), and application of 10% stretch on 20% GelMA further increased α-SMA relative to the unstretched condition (p ≤0.001). In contrast, gTM cells displayed consistently elevated α-SMA across all conditions, with no significant pairwise differences for stiffness or stretch. Cross-cell comparisons confirmed that gTM cells expressed significantly higher α-SMA than nTM cells at all matched conditions (all p ≤ 0.001; **Fig. S6-a**). Three-way ANOVA identified cell type as the sole significant main effect (F = 39.55, p < 0.0001), while hydrogel type (p =0.14), stretch type (p =0.25), and all interactions were non-significant.

**Fig. 6A.**
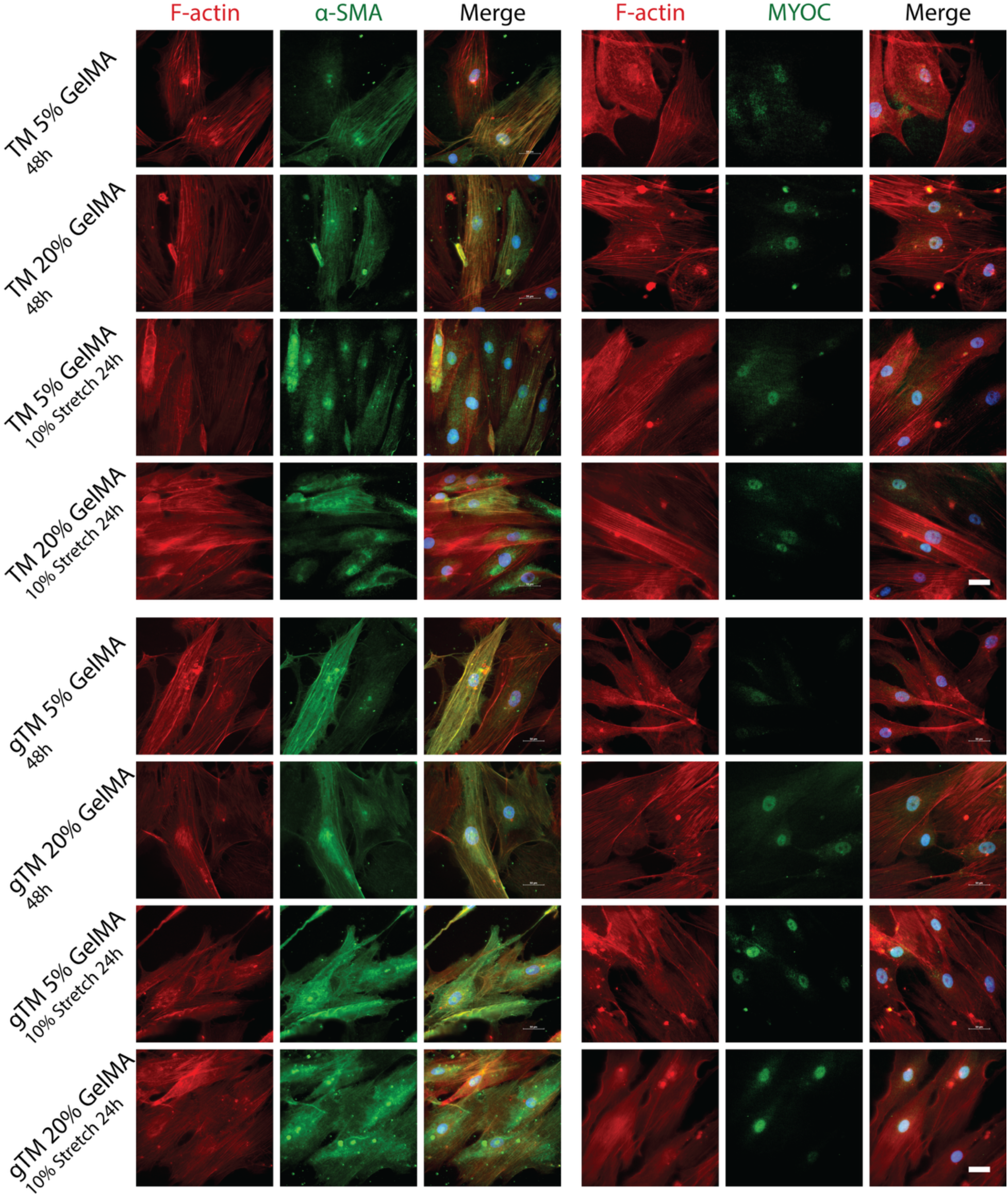
**Phenotypic Biomarkers of nTM and gTM Cells under Matrix Stiffness and Mechanical Stretch**. nTM and gTM cells were cultured on 5% and 20% GelMA hydrogels under static conditions for 48 h or with 10% stretch for 24 h following 24 h of cell stabilization. Representative immunofluorescence images show α-SMA and MYOC expression. Scale bar: 50 μm

**Fig. 6B.**
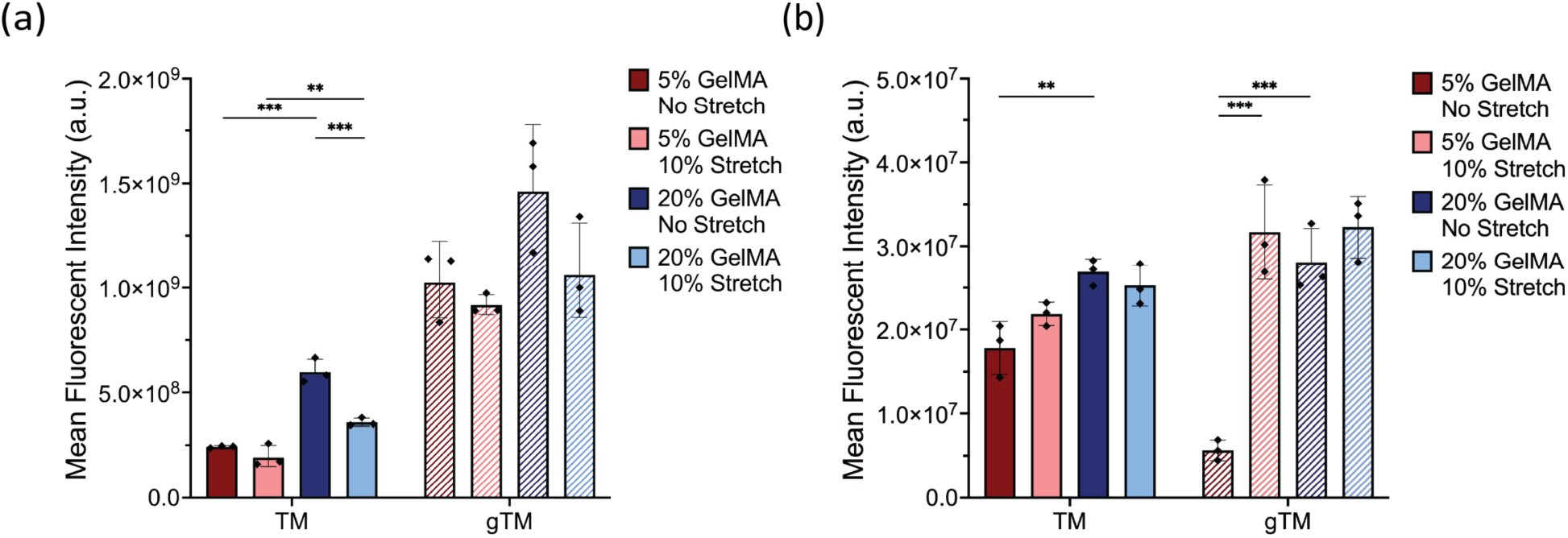
Quantitative Analysis of Phenotypic Biomarkers in nTM and gTM Cells under Varying Matrix Stiffness and Mechanical Stretch. Mean fluorescence intensity (a.u.) of α-SMA (a) and MYOC (b) in nTM and gTM cells within 5% and 20% GelMA hydrogels under no stretch or 10% stretch conditions. Within condition pairwise comparisons, α-SMA expression in nTM cells increased with substrate stiffness under both static and stretched conditions and was further elevated by 10% stretch on 20% GelMA, whereas gTM cells showed no significant pairwise differences across conditions. MYOC expression in nTM cells showed a modest but significant increase on 20% GelMA compared to 5% GelMA under no stretch conditions, while gTM cells exhibited a low baseline MYOC level on 5% GelMA without stretch that was strongly upregulated by either increased stiffness or the application of 10% stretch. Data are presented as mean ± SD with individual replicates shown (n = 3). Statistical significance: *p ≤ 0.05, **p ≤ 0.01, ***p ≤ 0.001.

**Fig 7A.**
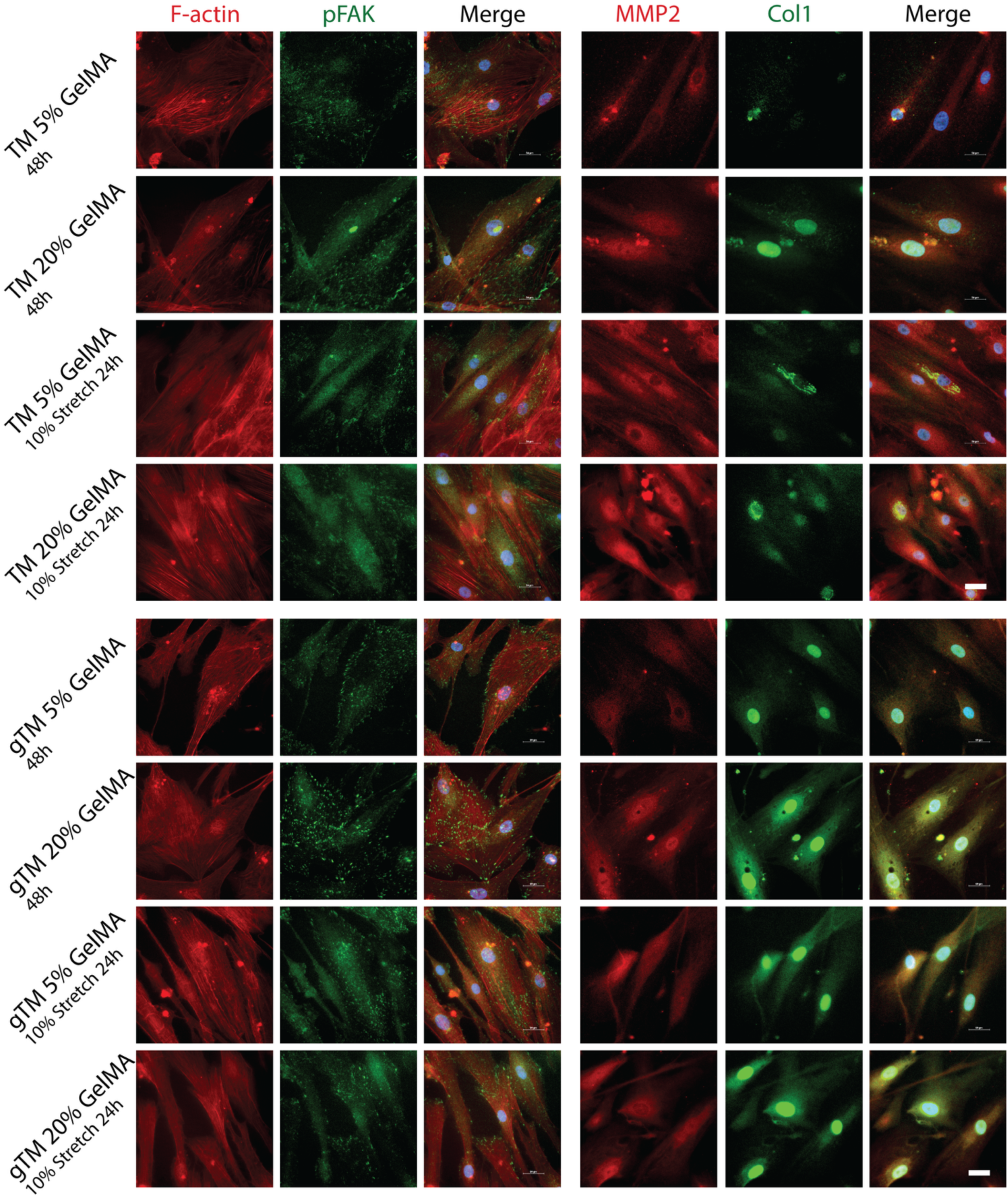
Focal Adhesion and ECM Biomarkers of nTM and gTM Cells under Matrix Stiffness and Mechanical Stretch. nTM and gTM cells were cultured on 5% and 20% GelMA hydrogels under static conditions for 48 h or with 10% static stretch for 24 h following 24 h of cell stabilization. Representative immunofluorescence images show pFAK, MMP2, and collagen I expression. Scale bar: 50 μm

**Fig 7B.**
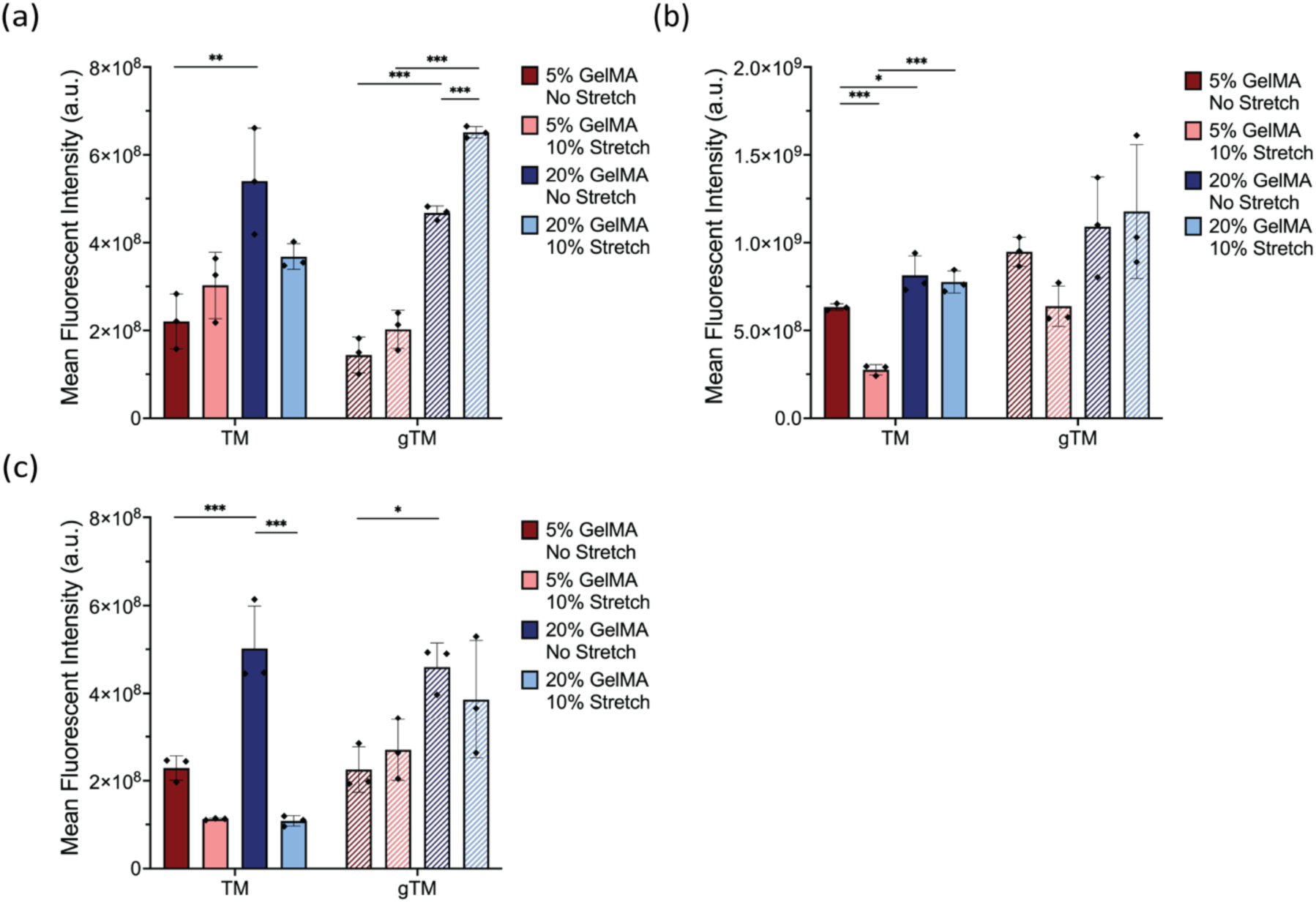
Quantitative Analysis of Focal Adhesion and ECM Turnover Biomarker of nTM and gTM Cells under various Matrix Stiffness and Mechanical Stretch. Quantitative analysis of (a) phosphorylated focal adhesion kinase (pFAK), (b) matrix metalloproteinase-2 (MMP2), and (c) type I collagen (COL1) expression, presented as mean fluorescence intensity in nTM and gTM cells within 5% and 20% GelMA hydrogels subjected to either no stretch or 10% stretch. pFAK expression was elevated by stiffness in both cell types; gTM cells showed a significant further increase with stretch on 20% GelMA (p ≤0.001), indicating enhanced mechanotransduction under combined conditions. In nTM cells, MMP2 was significantly reduced by stretch on compliant substrates (p ≤ 0.001), while COL1 was significantly reduced by stretch on stiff substrates (p ≤ 0.001). gTM cells showed no significant MMP2 changes across conditions; COL1 in gTM cells was modestly elevated by stiffness under static conditions (p ≤ 0.05) but was not significantly affected by stretch. Data are presented as mean ± SD with individual replicates shown (n = 3). Statistical significance: *p ≤ 0.05, **p ≤ 0.01, ***p ≤ 0.001.

**Fig. 6A** (right panels) and **Fig. S7** reveal a distinct pattern for MYOC expression that differed from α-SMA in both intensity and subcellular distribution. In nTM cells, MYOC signal was low on compliant substrates under static conditions and became more apparent on stiff substrates, where it appeared as a predominantly diffuse cytoplasmic and perinuclear distribution with only faint nuclear signal. Application of 10% stretch further increased MYOC signal on both 5% and 20% GelMA while retaining the similar diffuse cytoplasmic pattern. In gTM cells, MYOC was low on compliant substrates under static conditions but was markedly induced by application of either 10% stretch or increased substrate stiffness, remaining elevated under the combined stiff and stretched condition. Unlike nTM cells, gTM cells consistently exhibited a distinctly nuclear-localized MYOC distribution that co-localized with DAPI, with only modest cytoplasmic signal. **Fig. 6B-b** confirms these patterns quantitatively. In nTM cells, increasing substrate stiffness from 5% to 20% GelMA under static conditions elevated MYOC expression (p ≤0.01), while 10% stretch produced a trend toward higher MYOC on both substrates. In gTM cells, MYOC was strongly upregulated from the low baseline observed on 5% GelMA under static conditions: application of 10% stretch on 5% GelMA (p ≤0.001) and an increase in substrate stiffness to 20% GelMA under static conditions (p ≤0.001) each produced a pronounced increase in MYOC, with expression remaining elevated under the combined stiff and stretched condition. Cross-cell comparisons revealed that gTM cells expressed significantly lower MYOC than nTM cells on compliant substrates under static conditions (p ≤0.01) yet significantly higher MYOC than nTM cells under the combined 20% GelMA with 10% stretch condition (p ≤0.05) (**Fig. S6-b**). Both substrate stiffness and mechanical stretch modulated MYOC expression, with the strongest induction occurring in gTM cells as they transitioned from the compliant static baseline to either stiffer or stretched conditions. A three-way ANOVA further confirmed that substrate stiffness and mechanical stretch were the dominant factors driving MYOC expression, while cell type alone showed no significant main effect; instead, cell type influenced MYOC through significant interactions with both stiffness and stretch, including a significant three-way interaction, consistent with the amplified mechanical responsiveness observed in gTM cells.

### 2.6 Focal Adhesion and ECM Remodeling Responses in nTM and gTM Cells

pFAK, MMP2, and COL1 expression were quantified in nTM and gTM cells cultured on 5% and 20% GelMA hydrogels under no stretch or 10% equi-biaxial stretch. **Fig. 7A** (left panels) shows representative immunofluorescence images of pFAK expression. In nTM cells, pFAK signal was faint and diffusely distributed on compliant substrates, with punctate focal adhesion staining becoming more apparent on 20% GelMA. Application of 10% stretch modestly enhanced pFAK on stiff substrates. In gTM cells, pFAK signal was similarly low on compliant substrates but became markedly more intense on 20% GelMA, with prominent focal adhesion localization that was further enhanced under combined stiff and stretched conditions, producing the strongest pFAK signal observed across all groups. **Fig. 7B-a** quantifies these observations. Within nTM cells, increasing stiffness from 5% to 20% GelMA significantly elevated pFAK under no stretch conditions (p ≤ 0.01). In gTM cells, stiffness elevated pFAK under both no stretch (p ≤0.001) and 10% stretch (p ≤ 0.001) conditions, and application of 10% stretch on 20% GelMA further increased pFAK relative to the no stretch condition (p ≤ 0.001). Cross-cell-type comparison revealed that gTM cells expressed significantly higher pFAK than nTM cells only under the 20% GelMA 10% stretch condition (p ≤0.001; **Fig. S8-a**), suggesting that differences between cell types become most pronounced under combined stiff and stretched conditions. Three-way ANOVA identified hydrogel type (F =13.09, p =0.001) and stretch type (F =5.35, p =0.027) as significant main effects, while cell type (p = 0.68) and all interaction terms were non-significant.

**Fig. 7A** (right panels, red channel) reveals distinct MMP2 expression patterns between cell types. In nTM cells, MMP2 appeared as perinuclear and cytoplasmic staining that was low on compliant substrates under static conditions, increased visibly on stiff substrates, and was notably reduced by stretch on compliant substrates. In contrast, gTM cells displayed relatively uniform and low-level MMP2 signal across all conditions, with no apparent change in response to stiffness or stretch, visually consistent with a loss of MMP2 mechanoresponsiveness. **Fig. 7B-b** confirms these patterns. In nTM cells, stiffness significantly increased MMP2 under both no stretch (5% vs 20% GelMA; p ≤ 0.05) and 10% stretch (5% vs 20% GelMA; p ≤ 0.001) conditions. Conversely, 10% stretch on soft substrates significantly reduced MMP2 (5% GelMA no stretch vs 10% stretch; p ≤ 0.001). In contrast, gTM cells showed no significant pairwise differences across any conditions, indicating a loss of adaptive MMP2-mediated ECM remodeling responsiveness (**Fig. S8-b**). Three-way ANOVA identified both cell type (F = 8.25, p = 0.007) and hydrogel type (F = 9.04, p = 0.005) as significant main effects, while stretch type (p = 0.95) and all interactions were non-significant. **Fig. 7A** (right panels, green channel) shows COL1 expression. In nTM cells, COL1 signal was minimal on compliant substrates and substantially increased on 20% GelMA under static conditions, appearing as bright fibrillar and pericellular staining. Application of 10% stretch on stiff substrates markedly reduced COL1, restoring signal intensity closer to levels observed on compliant substrates. In gTM cells, COL1 was modestly elevated on stiff substrates under static conditions, but unlike nTM cells, stretch did not visibly diminish COL1 signal on 20% GelMA, resulting in persistently elevated staining under combined stiff and stretched conditions. **Fig. 7B-c** quantifies these trends. In nTM cells, increasing stiffness from 5% to 20% GelMA under no stretch conditions significantly elevated COL1 (p ≤0.001), while application of 10% stretch on 20% GelMA significantly reduced COL1 relative to the no stretch condition (p ≤0.001). Within gTM cells, stiffness modestly increased COL1 under no stretch conditions (5% vs 20% GelMA; p ≤0.05). Cross-cell-type comparison revealed significantly higher COL1 in gTM relative to nTM only under the 20% GelMA 10% stretch condition (p ≤0.01; **Fig. S8-c**), mirroring the pattern observed for pFAK. Three-way ANOVA confirmed cell type (F =18.86, p =0.0001) and hydrogel type (F =9.74, p =0.004) as significant main effects. Stretch type approached significance (p =0.052), and notably, the interaction between hydrogel type and stretch type was significant (F =8.10, p =0.008), indicating that the effect of stretch on COL1 expression depended on substrate stiffness. All other interactions were non-significant.

## 3. Discussion

The hydrogel-integrated microfluidic platform presented here overcomes a longstanding barrier in TM mechanobiology: the inability to apply coupled substrate stiffness and mechanical stretch within a single experimental system. Unlike previous systems restricted to uniaxial or biaxial deformation on fixed-stiffness substrates ^[20b]^, equi-biaxial hydraulic actuation of tunable GelMA hydrogels recapitulates the complex loading conditions experienced by nTM cells in vivo ^[3b, 23]^. The strain capacities achieved (**Fig. 3**) substantially exceed the stretch amplitudes commonly used in TM in vitro studies ^[8b, 9b]^, providing a broad operational window for modeling both normal and pathological conditions. FEA validation (**Fig. 4**) confirms the mechanical fidelity of the system and establishes a computational framework for rational design of future experiments.

FEA further revealed that the difference in material properties between the hyperelastic PDMS and GelMA hydrogel produces non-uniform stress redistribution across the composite construct. The substantial reduction in effective stress induced by GelMA (**Fig. 4D**) is functionally significant because it brings the mechanical environment into a more physiologically relevant range, allowing substrate stiffness rather than nonphysiological stress to serve as the primary mechanical variable sensed by cells. In the PDMS/5% GelMA composite, effective surface stress was reduced by ∼40% at the Top and Mid regions and ∼20% at the Edge, compared to ∼20% and ∼30%, respectively, in the PDMS/20% GelMA composite (**Fig. S3**). These findings indicate that softer hydrogels absorb a greater proportion of the transmitted mechanical load, effectively functioning as a tunable mechanical filter that governs both the magnitude and spatial distribution of stress experienced by cells. Because the GelMA was modeled as a biphasic material, fluid redistribution within the porous matrix under sustained static loading may further reduce the effective solid strain experienced by cells relative to the applied geometric membrane strain, particularly on softer 5% GelMA where higher water content allows greater fluid-phase load sharing. This poroelastic behavior reinforces substrate stiffness as a modulator of the local mechanical environment sensed by cells, even under nominally identical stretch conditions.

Beyond this material-dependent attenuation, the local balance between hoop and meridional stress varies across the dome, creating three mechanically distinct zones within a single construct (**Fig. 4C and 4D**): an equi-biaxial apex, a hoop-dominant mid-slope, and a highly anisotropic edge^[21]^. This spatial stress gradient allows region-specific cell responses to be examined within a single chip. These zones can be exploited to probe region-dependent cell responses without requiring additional chips. The morphological data directly validate this spatially graded environment. Region-dependent cell elongation in both TM cells on compliant substrates (Top vs. Edge: p ≤0.001; **Fig. 5B**) mirrors the stress anisotropy predicted by FEA, with equi-biaxial stress at the apex providing no preferential alignment cue while the progressively diverging hoop-to-meridional ratio drives increasing elongation and directional alignment at the mid-slope and edge. Cell orientation analysis confirmed this pattern: cells at the Top displayed random orientation, mid-slope cells aligned preferentially along the hoop stress direction, and edge cells aligned along the meridional direction toward the dome center (**Fig. 5A, S5**). The stronger significance of regional aspect ratio differences on compliant versus stiff substrates (p ≤0.001 vs. p ≤ 0.05) indicates that softer matrices permit greater cytoskeletal reorganization, whereas higher crosslink density partially constrains remodeling.

Notably, gTM cells on compliant substrates lost significant regional aspect ratio differences entirely, suggesting impaired mechanosensing specifically on substrates mimicking healthy tissue stiffness. On stiff substrates, gTM cells retained regional elongation (Top vs. Edge: p ≤0.01) but exhibited a divergent spreading area response: whereas nTM cells increased their spreading area from Top to Edge on both substrate stiffnesses, gTM cells on compliant substrates showed a significant decrease in area from Top to Edge (**Fig. 5C**). Cross-cell comparisons further highlighted these differences. gTM cells were larger than nTM at the Top on both substrates (5% GelMA: p ≤0.01; 20% GelMA: p ≤0.001), but this relationship reversed at the Edge on compliant substrates where nTM cells were larger (p ≤ 0.05). This pattern of elongation without proportional area expansion in gTM cells points toward cytoskeletal reorganization that lacks functional mechanoadaptation, a feature consistent with glaucomatous transformation ^[24]^.

Upstream of these morphological changes, pFAK reveals how focal adhesion signaling differentiates normal from glaucomatous mechanotransduction. Both cell types activated pFAK in response to stiffness, but gTM cells showed an enhanced response specifically under combined conditions (**Fig. 7B-a** **and S7-a**). Three-way ANOVA confirmed that the mechanical environment rather than cell type drove pFAK with hydrogel type (p = 0.001) and stretch type (p = 0.027) as significant main effects and cell type non-significant (p = 0.68). This pattern is consistent with altered cell mechanics and focal adhesion remodeling reported in glaucomatous tissue ^[10a, 24b]^ and suggests a condition-specific gain of function in focal adhesion signaling rather than a loss of mechanosensing, a shift that may have implications for downstream pathological signaling.

Despite this shared upstream sensitivity, α-SMA exhibited fundamentally different regulatory logic. nTM cells dynamically increased α-SMA with stiffness and stretch, whereas gTM cells maintained constitutively elevated expression irrespective of mechanical context, with cell type as the sole ANOVA driver (p <0.0001; **Fig. 6B-a, S6-a**). The dissociation between mechanosensitive pFAK and mechanically uncoupled α-SMA in gTM cells suggests that myofibroblast transformation, once consolidated, may operate as a stable phenotypic state ^[11a, 25]^ that persists independently of acute mechanical cues. The progressive α-SMA increase observed in nTM cells across conditions may represent the early, still-reversible phase of this transition, suggesting that early intervention during this responsive phase may warrant further investigation. MYOC regulation in TM cells appears to depend on both loading mode and baseline mechanical context. While 10% mechanical stretch upregulates MYOC mRNA in human TM cells^[12]^ and 24-hour sustained stretch increases extracellular myocilin secretion in porcine TM cells alongside heat shock and cytokine stress ^[26]^, no significant MYOC change is observed under 6% cyclic stretch^[17]^. These findings suggest that MYOC induction preferentially tracks quasi-static or sustained rather than oscillatory mechanical input. Consistent with this pattern, MYOC expression in our study was significantly elevated by increased substrate stiffness in both nTM and gTM cells. Quasi-static stretch further elevated MYOC in gTM cells on compliant substrates (p ≤ 0.001) but produced no additional significant increase on stiff substrates where MYOC was already elevated by stiffness, and only a non-significant upward trend in nTM cells (Fig. 6B-b, S6-b). This context-dependent stretch response, most pronounced when baseline MYOC is low, is consistent with the poroelastic attenuation of effective solid strain on softer GelMA discussed above, and suggests that MYOC responds to stretch primarily when stiffness-driven induction has not already saturated the response.

In nTM cells, mechanical stimulation produced a modest elevation of MYOC that remained distributed in a diffuse cytoplasmic and perinuclear pattern, whereas in gTM cells MYOC was induced far more strongly from a low compliant-static baseline and accumulated in a distinctly nuclear-localized distribution. This inducibility of MYOC in gTM cells, together with its re-localization to the nucleus, indicates that gTM cells are not mechanosensitively attenuated with respect to MYOC but rather exhibit amplified and qualitatively altered responses to mechanical cues. Notably, gTM cells expressed lower MYOC than nTM cells on compliant substrates under static conditions yet exceeded nTM cells under combined stiff and stretched conditions, a cross-cell reversal that is visible only under coupled mechanical stimulation. These observations suggest that MYOC regulation in gTM cells is decoupled from that of α-SMA, which remains constitutively elevated, and is instead governed by pathways that retain strong mechanical responsiveness ^[24a, 27]^. MYOC regulation may therefore remain amenable to mechanical modulation even in glaucomatous cells after other aspects of the disease phenotype have stabilized, a hypothesis that warrants further investigation.

The ECM markers further illustrate this selective pathway dysregulation. nTM cells dynamically regulated both MMP2 and COL1 in a context-dependent manner, while gTM cells lost MMP2 mechanoresponsiveness entirely yet retained modest stiffness-driven COL1 elevation (**Fig. 7B-b** **and -c**). The significant interaction between hydrogel type and stretch type for COL1 (p = 0.008) confirms that the effect of stretch on COL1 depends on substrate stiffness, and the absence of stretch-mediated COL1 suppression in gTM cells resulted in significantly higher collagen levels under combined stiff and stretched conditions (**Fig. S8-c**). This selective loss of matrix degradation coupled with partial retention of collagen deposition creates a net imbalance favoring ECM accumulation ^[28]^. Because stiffness itself promotes collagen production in gTM cells, the resulting mechanical environment may perpetuate a cycle of progressive matrix accumulation, while the capacity to counteract this buildup through MMP2-mediated degradation appears diminished.

The present findings demonstrate that gTM dysfunction does not reflect a uniform loss of mechanosensitivity but rather a selective dysregulation of distinct mechanotransduction and ECM remodeling pathways. While focal adhesion signaling remains mechanically responsive, with pFAK showing hyperresponsiveness under combined conditions, and MYOC is upregulated by both stiffness and stretch and accumulates in the nuclei of gTM cells, α-SMA has decoupled from mechanical cues entirely, and the capacity for MMP2-mediated matrix turnover and stretch-dependent COL1 suppression is impaired. This combination of amplified stiffness sensing, sustained collagen deposition, and reduced matrix degradation may contribute to a cycle of progressive tissue stiffening and outflow dysfunction, although direct causal relationships were not tested in this study ^[28]^. Importantly, several of these divergences between normal and glaucomatous cells were detectable only under combined stiff and stretched conditions, reinforcing the necessity of multiparametric mechanical platforms for resolving disease-relevant mechanobiological differences that remain obscured in single-parameter systems.

Our findings align with recent evidence that gTM cells possess intrinsically distinct mechanobiological programs. Direct measurements of TM cell traction have shown that gTM cells generate substantially larger forces than nTM cells across 1.5–80 kPa substrates (overlapping our range), with stress fiber density peaking at intermediate stiffness before declining^[29]^, paralleling the non-monotonic α-SMA and pFAK responses observed here. Mechanistically, stiffness- and TGFβ2-driven YAP/TAZ nuclear translocation has been linked to focal adhesion and actin remodeling via ERK/ROCK signaling^[30]^, a pathway consistent with our coupled pFAK, F-actin, and MMP2 findings. The recently quantified actomyosin-dominated cytoskeletal hierarchy in TM cells^[31]^ further rationalizes why our stretch responses track focal adhesion remodeling.

Several limitations should be noted. This study employed quasi-static stretch rather than the dynamic loading that mimics IOP fluctuations, which may elicit distinct response patterns^[16]^. The hydrogel stiffness range (1.23–21.47 kPa) spans lower-to-moderate disease progression and falls below values reported for advanced glaucomatous eyes^[3a, 4a]^, so additional crosslinking will be required for advanced disease modeling. The use of a single donor per cell type, consistent with prior primary TM cell studies, limits generalizability, and the 15-year age gap between donors (53 vs. 79 years) confounds disease status with donor age; age-matched multi-donor replication is needed for definitive attribution to glaucoma. Phenotypic drift in primary culture is also worth noting, since monolayer TM cells reportedly express MYOC at lower levels than fresh tissue and respond to dynamic stimulation^[12]^, and glaucomatous features such as CLANs may show variability across passages^[32]^. Our passage-3–5 measurements may thus capture an attenuated version of the native gTM response. Complementary analyses, including Western blot validation, gene expression profiling, multi-donor replication, and cyclic stretch studies, would further strengthen these observations and are warranted in future work.

## 4. Conclusion

This study presents a novel hydrogel-integrated microfluidic system that successfully addresses critical limitations in TM mechanobiology research while advancing lab-on-a-chip technology. Unlike conventional approaches, our GelMA-PDMS platform can simultaneously modulate substrate stiffness and apply physiologically relevant equi-biaxial tensile stress, enabling unprecedented control over cellular mechanical microenvironments. The integration of tunable GelMA hydrogels successfully recapitulated the stiffness range associated with glaucoma progression, while optimized surface modification protocols ensured robust hydrogel-PDMS integration that withstood up to 40% linear strain without mechanical failure. The close agreement between finite element models and experiments supports system reliability and provides a framework for rational design of future mechanobiological platforms.

Our data show that TM cells exhibit dramatically different mechanotransduction responses when exposed to coupled mechanical stimuli that occur physiologically. Most significantly, we demonstrate that disease-associated substrate stiffness and mechanical stretch converge to produce cellular responses distinct from those elicited by either stimulus alone, consistent with a pattern in which tissue stiffening promotes cellular behaviors associated with further ECM accumulation. The distinct mechanobiological profile of gTM cells, evidenced by stiffness-dependent pFAK activation and constitutively elevated α-SMA, provides new insights into how elevated intraocular pressure perpetuates disease progression through mechanotransduction pathways. This differential regulation of focal adhesion signaling, contractile markers, and ECM remodeling proteins under pathological stiffness conditions highlights candidate mechanistic targets for further study.

Notably, both increased substrate stiffness and 10% stretch (on compliant substrates) significantly upregulated MYOC expression in gTM cells, with pronounced nuclear localization, demonstrating that this disease-associated marker retains mechanosensitivity in gTM cells and suggesting a potential window for mechanical modulation of specific pathological features. This finding, combined with the platform’s ability to resolve condition-specific differences, supports further investigation of mechanotransduction-targeted interventions, including antifibrotic agents, as therapeutic strategies beyond traditional IOP reduction. From a technological perspective, our platform addresses a critical need in the organ chip and mechanobiology community for systems capable of applying complex, physiologically relevant mechanical stimuli while maintaining precise experimental control. The successful integration of multiple mechanical stimulation modalities into a single, optically transparent platform demonstrates the potential to investigate mechanobiology in other mechanically active tissues, including the cardiovascular, musculoskeletal, and respiratory systems. Future directions include investigations of cyclic mechanical loading, integration of flow-based shear stress, expansion to three-dimensional co-culture systems, and therapeutic screening. This hydrogel-integrated microfluidic system provides a versatile platform for mechanistic investigation that bridges the gap between simplified in vitro models and complex in vivo mechanical environments, providing both technological tools and biological insights that advance understanding of mechanobiology while supporting development of targeted therapeutic strategies for glaucoma and other mechanically mediated diseases.

## 5. Materials and Methods

### 5.1 Hydraulically Actuated Microfluidic System

A hydrogel-integrated microfluidic chip was designed to enable simultaneous testing of four conditions under precise mechanical control. The inflatable microfluidic chip was designed in AutoCAD (Autodesk, USA) and fabricated on microscope slides (Fisherbrand 12-550-A3, Fisher Scientific, USA) using five PDMS-based layers (Fig. 1A): (1) hydraulic pressure channel layer, (2) PDMS membrane layer, (3) hydrogel channel layer, (4) well layer, and (5) port layer. The hydraulic pressure channel, hydrogel channel, and well layers were fabricated from 250 μm-thick PDMS sheets (BISCO HT-6240, Rogers Corp., USA) using a vinyl cutter (CAMM-1 GX-24 24“, Roland DGA Corp., USA). The hydrogel channel layer contained 4 mm-diameter hydrogel regions connected by 0.6 mm-wide microfluidic channels. A 0.1 mm-thick PDMS membrane (GASKET-UT-100, SiMPore Inc., USA) was used to ensure uniform membrane properties across experiments. The port layer was cast from Sylgard 184 PDMS (Dow Inc., USA) mixed at a 10:1 base-to-curing-agent ratio, and inlet/outlet holes were created using a 1 mm biopsy punch.

Before assembly, glass slides were cleaned sequentially with acetone, isopropyl alcohol (IPA), and deionized water. The five layers were then sequentially bonded onto the glass substrate by oxygen plasma treatment (PE-25, Plasma Etch Inc., USA; 100 W, 30 s, 200 mTorr O₂). After assembly, the chip was cured overnight at 70°C to strengthen interlayer bonding. The completed chip was sterilized with 70% ethanol, rinsed with deionized water, and exposed to UV light (254 nm) for 30 min. Quality control measurements confirmed dimensional tolerances within ±5% and chip reproducibility with a coefficient of variation below 10%.

### 5.2 Hydrogel Preparation and Integration

#### GelMA precursor solution preparation

GelMA precursor solutions were prepared at final concentrations of 5%, 10%, and 20% (w/v) by dissolving PhotoGel® 50% DS in photoinitiator-containing phosphate-buffered saline (PBS, pH 7.4). Lithium phenyl-2,4,6-trimethylbenzoylphosphinate (LAP) served as the photoinitiator for 365 nm photocrosslinking^[33]^. A 0.5% (w/v) LAP stock solution was prepared in PBS, vortex-mixed at room temperature until fully dissolved, and sterile-filtered through a 0.22 μm syringe filter. To achieve the target GelMA concentrations, 10, 5, and 2.5 mL of the LAP stock solution were added directly to individual PhotoGel® 50% DS stock bottles, yielding 5%, 10%, and 20% GelMA, respectively. Each mixture was stirred at 100 rpm and 40 °C for 1 h using a magnetic stir bar until homogeneous, then maintained at 40 °C until use to prevent physical gelation.

#### Chip surface functionalization

To enable covalent anchoring of GelMA to the PDMS hydrogel channel layer, the assembled chip was surface-functionalized with 3-(trimethoxysilyl)propyl methacrylate (TMSPMA; M6514, Sigma-Aldrich, USA) as described in **Fig.2** ^[34]^. The chip was first exposed to oxygen plasma (100 W, 60 s, 200 mTorr O₂) to generate surface hydroxyl groups on the PDMS. Immediately after plasma treatment, the chip was incubated with 10% (v/v) TMSPMA in ethanol/deionized water (1:1, v/v) at 50 °C for 1 h, rinsed thoroughly with deionized water to remove unreacted silane, and dried under UV light (254 nm) for 1 h.

#### Sacrificial coverslip preparation

Glass coverslips (8 mm diameter) used to temporarily seal the hydrogel channel during GelMA loading were prepared in parallel. Coverslips were sequentially cleaned with acetone, isopropyl alcohol, and deionized water, then treated with oxygen plasma (100 W, 60 s, 200 mTorr O₂). To render their surfaces non-adhesive and allow clean release from the crosslinked hydrogel, the coverslips were coated with Sigmacote® (SL2, Sigma-Aldrich, USA) overnight at 50 °C, rinsed with deionized water, and dried under UV light for 1 h.

#### Hydrogel loading and photocrosslinking

Prior to hydrogel loading, deionized water was introduced into the hydraulic pressure channel to flatten the PDMS membrane and establish a zero-pressure reference state. A patterned Rubylith® photomask (RU3, Ulano Corp., USA) was aligned beneath the chip to define four circular crosslinking regions registered to the hydrogel wells. The Sigmacote-treated coverslips were then placed over the hydrogel channel layer, non-adhesive side facing downward, to form temporarily sealed microfluidic channels. Preheated GelMA precursor solution (40 °C) was introduced through the center inlet by gentle pipetting, which drove uniform filling of all four wells while minimizing bubble entrapment. The hydrogel-filled chip was exposed to 365 nm UV light at a total dose of 8 J/cm² through the photomask to selectively crosslink GelMA within the masked well regions ^[33b]^. After crosslinking, the chip was transferred to a cooling plate at 4 °C for 5 min to facilitate coverslip release. The coverslips were then lifted off, uncrosslinked GelMA in the channel regions outside the masked wells was aspirated, and the crosslinked hydrogel surfaces were immediately submerged in PBS and kept hydrated until cell seeding.

#### Optimization of UV exposure and mechanical characterization

To identify crosslinking conditions that produced dimensionally stable 5% GelMA hydrogels, samples were exposed to 365 nm UV light for 60, 90, 120, 150, 180, or 300 s. Following crosslinking, samples were cooled for 5 min and incubated in PBS (pH 7.4) at 37 °C for 24 h. Hydrogel shrinkage was quantified by imaging the projected surface area using dye-enhanced PBS for contrast, and comparing post-incubation to pre-incubation dimensions. The Young’s modulus of each GelMA formulation was determined by shear rheology. Storage and loss moduli were recorded during an amplitude sweep from 0 to 200% strain at 1 Hz using a 25 mm parallel-plate geometry. Hydrogels were equilibrated in PBS at room temperature for 20 min prior to testing, and excess surface PBS was blotted away immediately before measurement.

### 5.3 Mechanical Characterization

The mechanical testing apparatus consisted of a 1 mL BD™ reusable syringe connected via plastic barbed tube fittings and polyurethane tubing to the hydrogel-integrated microfluidic system. The syringe was actuated using a precision syringe pump. Barbed fittings were inserted into inlets with a slight interference fit to create pressure-tight seals. Prior to each experiment, a constant flow was applied to remove air bubbles from the hydraulic pressure channels and to level the PDMS membrane by confirming zero internal pressure. Following membrane leveling, a plastic sliding clamp was applied to the outlet tubing to prevent leakage during testing. All experiments were conducted at room temperature.

Optical measurements were performed using an optical microscope equipped with a near-infrared enhanced CMOS camera. Image analysis was conducted using open-source software (Fiji/ImageJ2, NIH). Membrane strain was calculated assuming semicircular deflection geometry, with linear strain (ε) determined from the change in deflected arc length (ΔL) and original membrane diameter (L₀). The detailed equation is provided in the SI.

Finite element analysis (FEA) was performed using FEBio to validate experimental results ^[35]^. We modeled three membrane configurations: PDMS alone, a PDMS/5% GelMA composite, and a PDMS/20% GelMA composite. The PDMS membrane alone (disc: 2 mm radius, 0.1 mm height) was modeled with Mooney–Rivlin hyperelastic constitutive behavior. For the composite configurations, a GelMA hydrogel layer (2 mm radius, 0.25 mm height) was added above the PDMS membrane; this layer was modeled using a biphasic formulation combining Veronda–Westmann hyperelastic behavior for the solid phase and water for the fluid phase^[22]^. We determined the constitutive constants (C_1_, C_2_) by iteratively fitting simulated dome deflections to experimental pressure–deflection data. For all configurations, boundary conditions included zero-displacement constraints at the sidewalls of both the PDMS and hydrogel layers, with hydraulic pressure applied to the bottom surface of the PDMS membrane. This FEA framework enabled predictive design and validation of membrane mechanics across varying hydrogel formulations.

### 5.4 Biological Characterization

Normal TM cells were isolated from de-identified postmortem human donor eyes (53-year-old male, no glaucoma history). gTM cells were obtained from a 79-year-old male donor with documented open angle glaucoma and associated ocular conditions (Utah Lions Eye Bank). All tissues were obtained in a de-identified manner, and no personally identifiable information was available to the investigators. The use of human donor tissue was determined to be exempt from Institutional Review Board oversight. Tissue procurement by the eye bank complied with informed consent procedures and adhered to the tenets of the Declaration of Helsinki.

Primary nTM and gTM cells were isolated from juxtacanalicular and corneoscleral meshwork regions. Both cell types were cultured in collagen type I-coated (Advanced Biomatrix #5005, USA) T75 flasks at 10,000 cells/cm² seeding density under standard conditions (37°C, 5% CO₂). Passages 3-5 were used for all experiments, cultured in TM Cell Medium (TMCM) (#6591, ScienCell Research Laboratories Inc., USA). Cells were passaged at 80-90% confluency using TrypLE™ Express Enzyme (1×), phenol red-free (#12604013, Thermo Fisher Scientific Inc., USA). Culture medium was exchanged daily following cell seeding on GelMA hydrogels. For static culture conditions, cells were cultured for 48 h post-seeding. For mechanical stimulation experiments, 10% stretch was applied for 24h following an initial 24h adaptation period on hydrogels.

Cell viability was assessed using CellTracker™ Green CMFDA fluorescent probes (C7025, Invitrogen Co., USA). CellTracker™ was diluted 1:1000 in TMCM. The existing culture medium was removed and replaced with CellTracker-containing medium, followed by a 30-minute incubation at 37°C. After incubation, the staining solution was replaced with fresh TMCM, and cells were imaged using a Nikon Eclipse Ti-E inverted microscope with a FITC filter (excitation/emission: 492/517 nm). Cells were washed once with PBS, then fixed with 4% paraformaldehyde in PBS (J61899.AK, Thermo Fisher Scientific Inc., USA) for 30 min at room temperature. Following three 5-min PBS washes, cells were permeabilized with 0.3% Triton X-100 (X-100, Sigma-Aldrich, USA) for 15 min at room temperature. After three additional 5-min PBS washes, non-specific binding was blocked with 20% bovine serum albumin (BSA) (A8022, Sigma-Aldrich, USA) for 24 h at 4°C. Primary antibodies were applied overnight at 4°C following BSA removal: anti-α-smooth muscle actin [1A4] (ab7817, Abcam, UK) (1:100), myocilin [F-12] (sc-137233, Santa Cruz Biotechnology, USA) (1:100), anti-FAK (phosphor-Y397) [EP2160Y] (ab81298, Abcam, UK) (1:400), anti-MMP2 (ab97779, Abcam, UK) (1:100), and anti-collagen type I (COL1A1) (MAB3391, Sigma-Aldrich, USA) (1:100). Following three 5-min PBS washes, secondary antibodies were applied for 1 h at room temperature in darkness: phalloidin-TRITC (P1951, Sigma-Aldrich, USA) (1:1000), anti-rabbit Alexa Fluor 488 (ab150077, Abcam, UK) (1:200), anti-rabbit Alexa Fluor 594 (A-11012, ThermoFisher Scientific Inc., USA) (1:1000), and anti-mouse CF 488A (SAB4600388, Sigma-Aldrich, USA) (1:200). After three 5-min PBS washes, nuclei were counterstained with Hoechst 33258 (H3569, ThermoFisher Scientific Inc., USA) (1:3000) for 15 min at room temperature, followed by a final 5-min PBS wash. Samples were imaged using a Nikon Eclipse Ti-E inverted microscope.

### 5.5 Image Processing and Statistical Analysis

Initial image capture and processing were performed using Nikon NIS-Elements AR 5.0 (Nikon Corp., Japan) for large tile images, orthogonal z-stack views, and the addition of scale bars. All images were then processed and analyzed using MATLAB (MathWorks, USA). Images were taken mainly from three different locations (Top, Mid, and Edge) using the Cartesian coordinate system. Top images were taken from the center of the hydrogel and considered as (0, 0) in the Cartesian coordinate. Mid images were taken 2 mm to the right of the center, at (2, 0) in the Cartesian coordinate system. Edge images were taken 4 mm to the right of the center, at (4, 0) in the Cartesian coordinate system. These three locations correspond to the equi-biaxial apex, hoop-dominant mid-slope, and highly anisotropic edge zones identified by FEA.

For morphological analysis under 10% stretch, cell aspect ratio was calculated as the ratio of the minor axis to the major axis, where values approaching 1 indicate a circular morphology and lower values indicate greater elongation. Cell spreading area was measured from the segmented cell boundaries. Cell orientation was determined from the angle of the major axis relative to the hoop stress direction and presented as orientation distribution plots.

For immunofluorescence quantification, mean fluorescence intensity (arbitrary units, a.u.) was measured for each biomarker: α-SMA, MYOC, pFAK, MMP2, and COL1. Fluorescence intensity was quantified from images of nTM and gTM cells cultured on 5% and 20% GelMA hydrogels under no stretch or 10% stretch conditions.

Statistical analysis and graphing were performed using Prism (GraphPad Software, USA). Each data point represents an independent biological replicate from one well or hydrogel, with n ≥ 3 per condition, and all data are presented as mean ± standard deviation (SD). For each biomarker, a three-way analysis of variance (ANOVA) was conducted with cell type, hydrogel type, and stretch condition as fixed factors to evaluate main effects and interaction terms. The cell type factor compared nTM and gTM cells, the hydrogel type factor compared 5% and 20% GelMA, and the stretch condition factor compared no stretch with 10% stretch. Pairwise comparisons were then performed using Tukey’s post hoc test with multiplicity-adjusted p values. Within each cell type, pairwise comparisons assessed the effects of substrate stiffness and mechanical stretch (**Fig. 6B**, **7B**). Between cell types, matched stiffness and stretch conditions were compared to evaluate differences between nTM and gTM cells (**Fig. S6, S7**). Statistical significance was defined as p ≤ 0.05 and denoted as *p ≤ 0.05, **p ≤ 0.01, ***p ≤ 0.001.

## Acknowledgements

This work was supported by the Ascender Grant from the University of Utah and by the Department of Mechanical Engineering at the University of Utah. This work was also supported by the John A. Moran Foundation and an Unrestricted Grant from Research to Prevent Blindness to the Moran Eye Center at the University of Utah. Authors also acknowledge the members of the Biomedical Micro-NanoSystems Lab for their support and contributions. We thank Dr. Bates for granting access to the rheometer used to measure the material properties of the hydrogels.

## References

[1] Y.-C. Tham, X. Li, T. Y. Wong, H. A. Quigley, T. Aung, C.-Y. Cheng, Ophthalmology 2014, 121, 2081.

[2] a)N. Zhang, J. Wang, Y. Li, B. Jiang, Scientific Reports 2021, 11, 13762; b)R. N. Weinreb, T. Aung, F. A. Medeiros, JAMA 2014, 311, 1901.

[3] a)K. Wang, G. Li, A. T. Read, I. Navarro, A. K. Mitra, W. D. Stamer, T. Sulchek, C. R. Ethier, Scientific Reports 2018, 8, 5848; b)W. D. Stamer, T. S. Acott, Current Opinion in Ophthalmology 2012, 23.

[4] a)J. A. Last, T. Pan, Y. Ding, C. M. Reilly, K. Keller, T. S. Acott, M. P. Fautsch, C. J. Murphy, P. Russell, Investigative Ophthalmology & Visual Science 2011, 52, 2147; b)K. Wang, M. A. Johnstone, C. Xin, S. Song, S. Padilla, J. A. Vranka, T. S. Acott, K. Zhou, S. A. Schwaner, R. K. Wang, T. Sulchek, C. R. Ethier, Investigative Ophthalmology & Visual Science 2017, 58, 4809.

[5] a)J. A. Vranka, M. J. Kelley, T. S. Acott, K. E. Keller, Experimental Eye Research 2015, 133, 112; b)T. S. Acott, M. J. Kelley, Experimental Eye Research 2008, 86, 543.

[6] P. L. Kaufman, Experimental Eye Research 2020, 197, 108105.

[7] M. A. Johnstone, W. G. Grant, Am J Ophthalmol 1973, 75, 365.

[8] a)D. WuDunn, Experimental Eye Research 2009, 88, 718; b)P. B. Liton, P. Gonzalez, Journal of Glaucoma 2008, 17, 378.

[9] a)M. Bikuna-Izagirre, J. Aldazabal, L. Extramiana, J. Moreno-Montañés, E. Carnero, J. Paredes, Biotechnology and Bioengineering 2022, 119, 2698; b)V. Vittal, A. Rose, K. E. Gregory, M. J. Kelley, T. S. Acott, Investigative Ophthalmology & Visual Science 2005, 46, 2857.

[10] a)M. Lakk, D. Križaj, American Journal of Physiology-Cell Physiology 2021, 320, C1013; b)D. A. Ryskamp, A. M. Frye, T. T. Phuong, O. Yarishkin, A. O. Jo, Y. Xu, M. Lakk, A. Iuso, S. N. Redmon, B. Ambati, G. Hageman, G. D. Prestwich, K. Y. Torrejon, D. Krizaj, Sci Rep 2016, 6, 30583; c)O. Yarishkin, T. T. T. Phuong, J. M. Baumann, M. L. De Ieso, F. Vazquez-Chona, C. N. Rudzitis, C. Sundberg, M. Lakk, W. D. Stamer, D. Krizaj, J Physiol 2021, 599, 571.

[11] a)E. R. Tamm, A. Siegner, A. Baur, E. LÜTjen-Drecoll, Experimental Eye Research 1996, 62, 389; b)S. M. Thomasy, J. A. Wood, P. H. Kass, C. J. Murphy, P. Russell, Investigative Ophthalmology & Visual Science 2012, 53, 952.

[12] E. R. Tamm, P. Russell, D. L. Epstein, D. H. Johnson, J. Piatigorsky, Invest Ophthalmol Vis Sci 1999, 40, 2577.

[13] Y. Okada, T. Matsuo, H. Ohtsuki, Japanese Journal of Ophthalmology 1998, 42, 90.

[14] a)K. E. Keller, M. J. Kelley, T. S. Acott, Investigative Ophthalmology & Visual Science 2007, 48, 1164; b)D. L. Fleenor, A. R. Shepard, P. E. Hellberg, N. Jacobson, I.-H. Pang, A. F. Clark, Investigative Ophthalmology & Visual Science 2006, 47, 226.

[15] a)Y.-F. Yang, Y. Y. Sun, D. M. Peters, K. E. Keller, Frontiers in Cell and Developmental Biology 2022, Volume 10 - 2022; b)J. A. Wood, C. T. McKee, S. M. Thomasy, M. E. Fischer, N. M. Shah, C. J. Murphy, P. Russell, Investigative Ophthalmology & Visual Science 2011, 52, 9298.

[16] C. Luna, G. Li, P. B. Liton, D. L. Epstein, P. Gonzalez, Mol Vis 2009, 15, 534.

[17] M. Lakk, D. Krizaj, Am J Physiol Cell Physiol 2021, 320, C1013.

[18] V. K. Raghunathan, J. T. Morgan, B. Dreier, C. M. Reilly, S. M. Thomasy, J. A. Wood, I. Ly, B. C. Tuyen, M. Hughbanks, C. J. Murphy, P. Russell, Investigative Ophthalmology & Visual Science 2013, 54, 378.

[19] H. Zou, R. Yuan, Q. Zheng, Y. Huo, M. Lang, S. Ji, Z. Zheng, J. Ye, Cellular Physiology and Biochemistry 2014, 33, 1215.

[20] a)S. Cosson, M. P. Lutolf, Scientific Reports 2014, 4, 4462; b)S. An, S. Y. Han, S.-W. Cho, BioChip Journal 2019, 13, 306.

[21] M. Kim, K. Choi, D. Križaj, J. Kim, Nature Communications 2025, 16, 9944.

[22] D. R. Veronda, R. A. Westmann, Journal of Biomechanics 1970, 3, 111.

[23] D. R. Overby, W. D. Stamer, M. Johnson, Experimental Eye Research 2009, 88, 656.

[24] a)W. D. Stamer, A. F. Clark, Experimental Eye Research 2017, 158, 112; b)J. T. Morgan, V. K. Raghunathan, Y. R. Chang, C. J. Murphy, P. Russell, Oncotarget 2015, 6, 15362.

[25] A. R. Muralidharan, R. Maddala, N. P. Skiba, P. V. Rao, Invest Ophthalmol Vis Sci 2016, 57, 6482.

[26] A. M. Anderssohn, K. Cox, K. O’Malley, S. Dees, M. Hosseini, L. Boren, A. Wagner, J. M. Bradley, M. J. Kelley, T. S. Acott, Invest Ophthalmol Vis Sci 2011, 52, 7548.

[27] E. R. Tamm, P. Russell, D. L. Epstein, D. H. Johnson, J. Piatigorsky, Investigative Ophthalmology & Visual Science 1999, 40, 2577.

[28] a)K. E. Keller, D. M. Peters, Clin Exp Ophthalmol 2022, 50, 163; b)P. P. Pattabiraman, R. Maddala, P. V. Rao, J Cell Physiol 2014, 229, 927.

[29] A. Karimi, M. Aga, T. Khan, D. D’Costa S, S. Cardenas-Riumallo, M. Zelenitz, M. J. Kelley, T. S. Acott, Acta Biomater 2024, 175, 138.

[30] H. Li, V. Raghunathan, W. D. Stamer, P. S. Ganapathy, S. Herberg, Front Cell Dev Biol 2022, 10, 844342.

[31] A. Karimi, A. Stanik, H. Golchin, D. Fuller, M. Aga, E. White, M. J. Kelley, T. S. Acott, Acta Biomater 2025, 203, 478.

[32] M. Peng, N. P. Rayana, J. Dai, C. K. Sugali, H. Baidouri, A. Suresh, V. K. Raghunathan, W. Mao, Exp Eye Res 2022, 220, 109097.

[33] a)A. K. Nguyen, P. L. Goering, V. Reipa, R. J. Narayan, Biointerphases 2019, 14, 021007; b)S. Sharifi, H. Sharifi, A. Akbari, J. Chodosh, Scientific Reports 2021, 11, 23276.

[34] Z. Dong, Q. Yuan, K. Huang, W. Xu, G. Liu, Z. Gu, RSC Advances 2019, 9, 17737.

[35] S. A. Maas, B. J. Ellis, G. A. Ateshian, J. A. Weiss, Journal of Biomechanical Engineering 2012, 134.

