## Supplemental files for "Investigating the coupled effects of stiffness and stretch on the trabecular meshwork cells using a hydrogel-integrated microfluidic system"

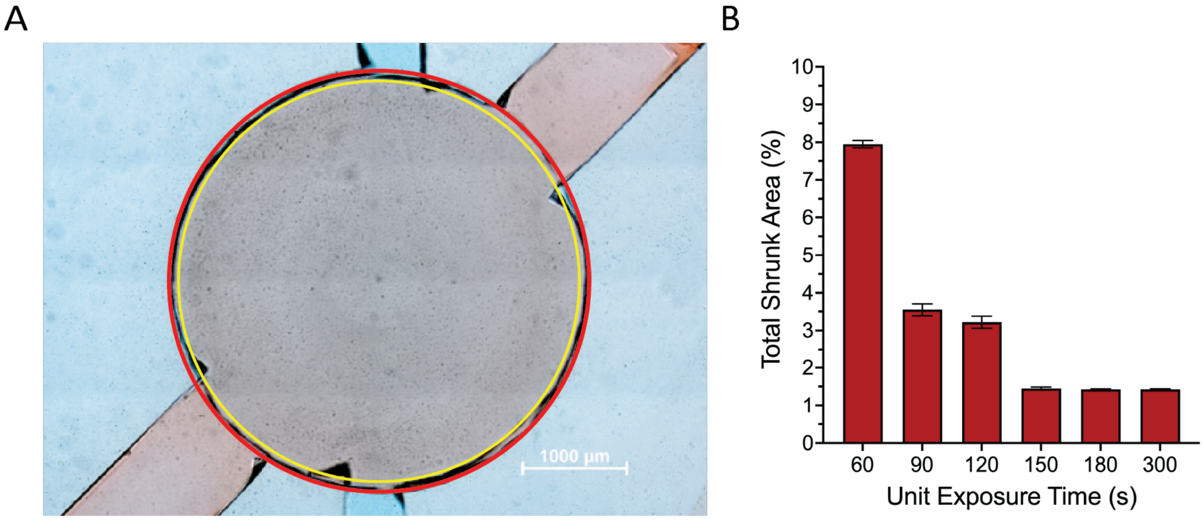


**Fig. S1.** Hydrogel Surface Area and UV Exposure Analysis (A) A photo of 5% GelMA hydrogel after exposure to UV for 150 seconds and incubated for 24h. The red line shows the original circumference of the hydrogel before incubation, and the yellow dotted line shows the shrunk circumference of the hydrogel after incubation for 24h. (B) Total shrunk area of 5% GelMA hydrogel based on UV exposure time after incubation for 24h. After UV exposure for 150 seconds, the total shrunk area was consistent after incubation for 24h.

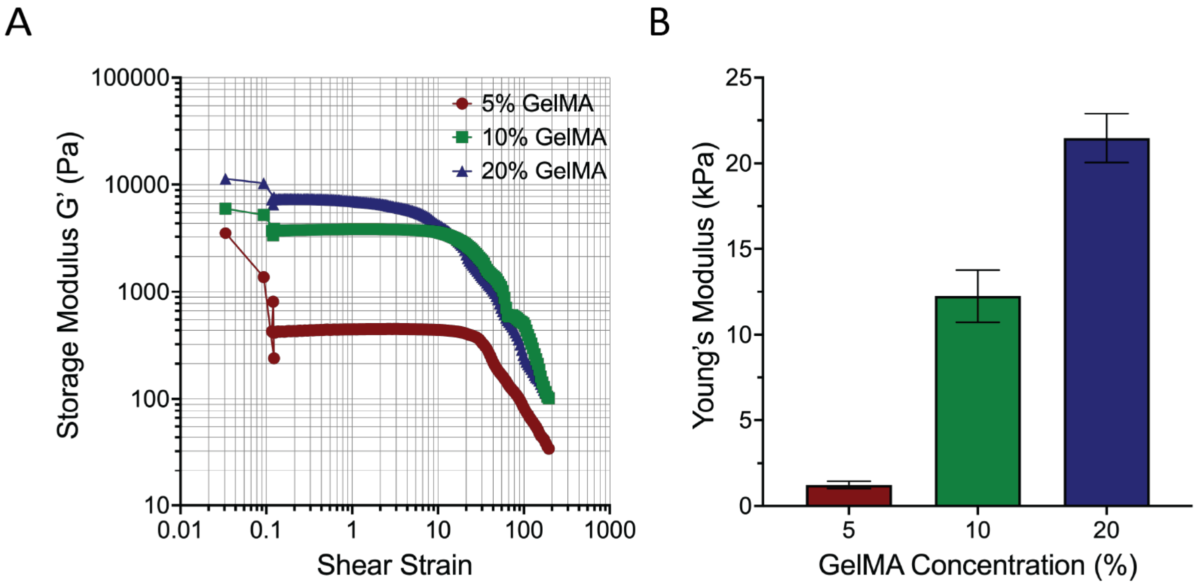


**Fig. S2.** Young's Modulus of GelMA hydrogel with different concentrations using a rheometer. (A) Representative data of each GelMA hydrogel. Storage modulus values were obtained by increasing the shear strain rate. (B) Young's modulus of GelMA hydrogel with different GelMA Concentrations. Each Young's modulus value was calculated based on the storage modulus values.

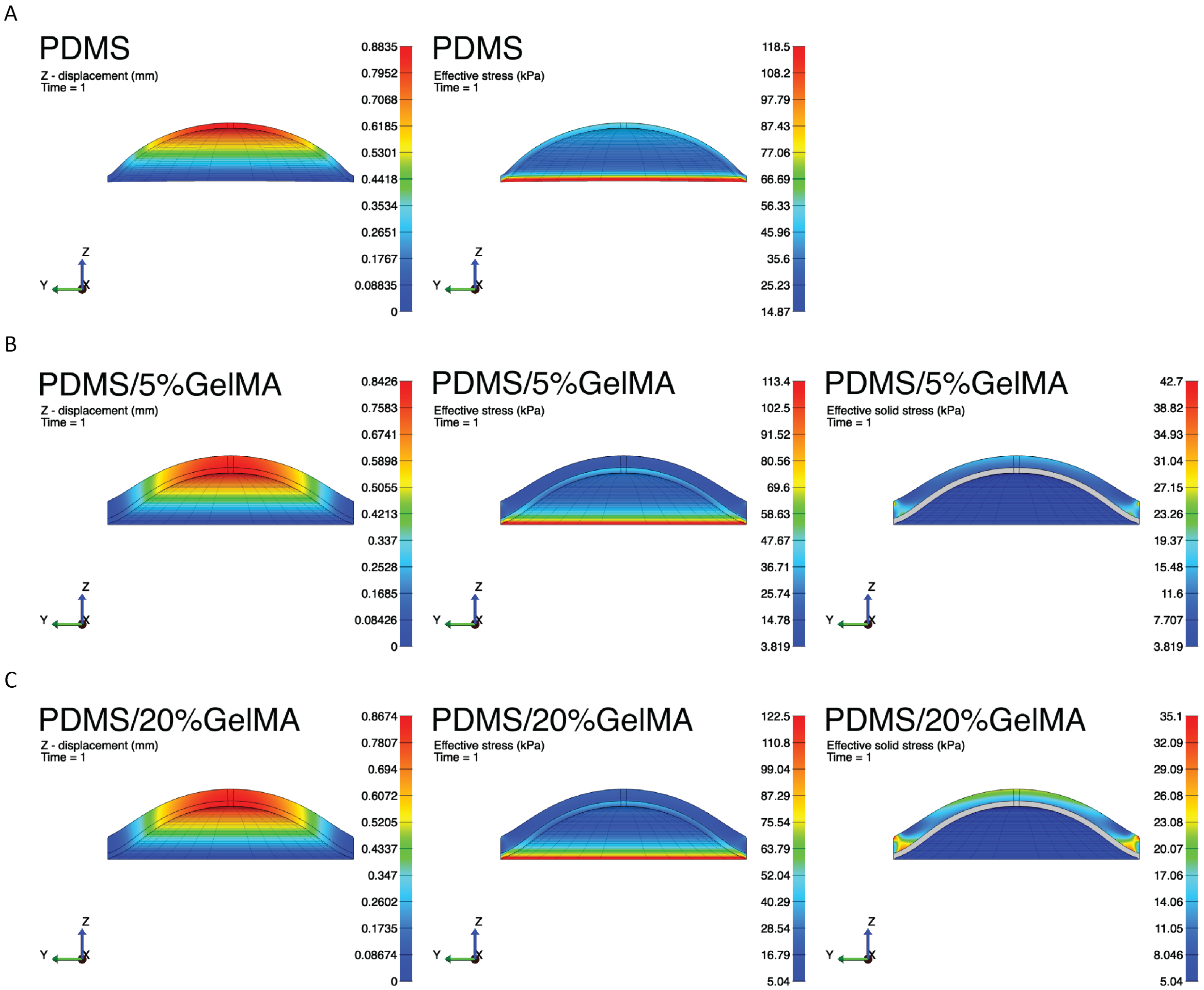


**Fig. S3**. Finite Element Method (FEM) analysis of membrane deflection and stress under various pressure for three cases (PDMS only, 5% GelMA on PDMS, and 20% GelMA on PDMS). (A) Cross-sectional views illustrating the displacement field and effective stress of the PDMS membrane. (B) Cross-sectional views illustrating the displacement field, effective stress, and effective solid stress of the 5% GelMA on PDMS membrane. (C) Cross-sectional views illustrating the displacement field, effective stress, and effective solid stress of the 20% GelMA on PDMS membrane. Color scales indicate the magnitude of displacements (mm) and the stress levels of each case in (kPa). Effective solid stress isolates the stress gradient within the hydrogel enabling visualization of the stress gradient distribution on the cell-bearing surface.

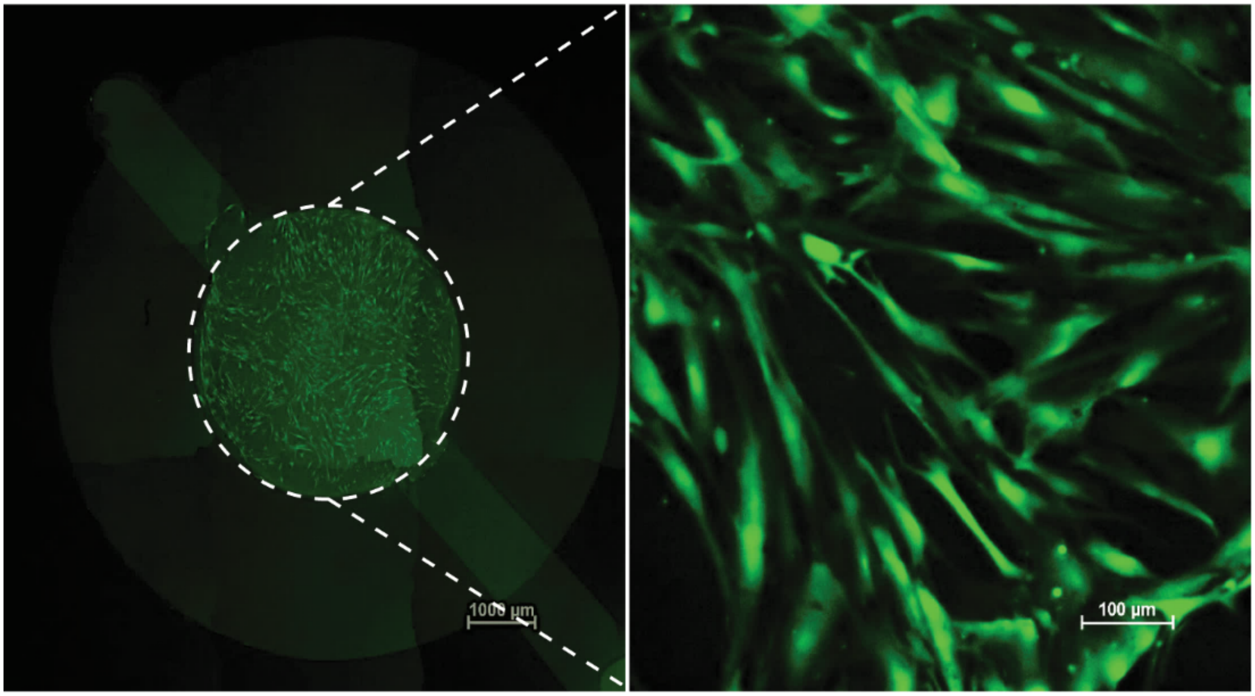


**Fig. S4.** Cell Viability Test on GelMA Hydrogel. (A) Large tile image of trabecular meshwork cells on the hydrogel in the well (B) Close-up image of trabecular meshwork cells on the hydrogel. Trabecular meshwork cells were only seeded and proliferated on the hydrogel surface.

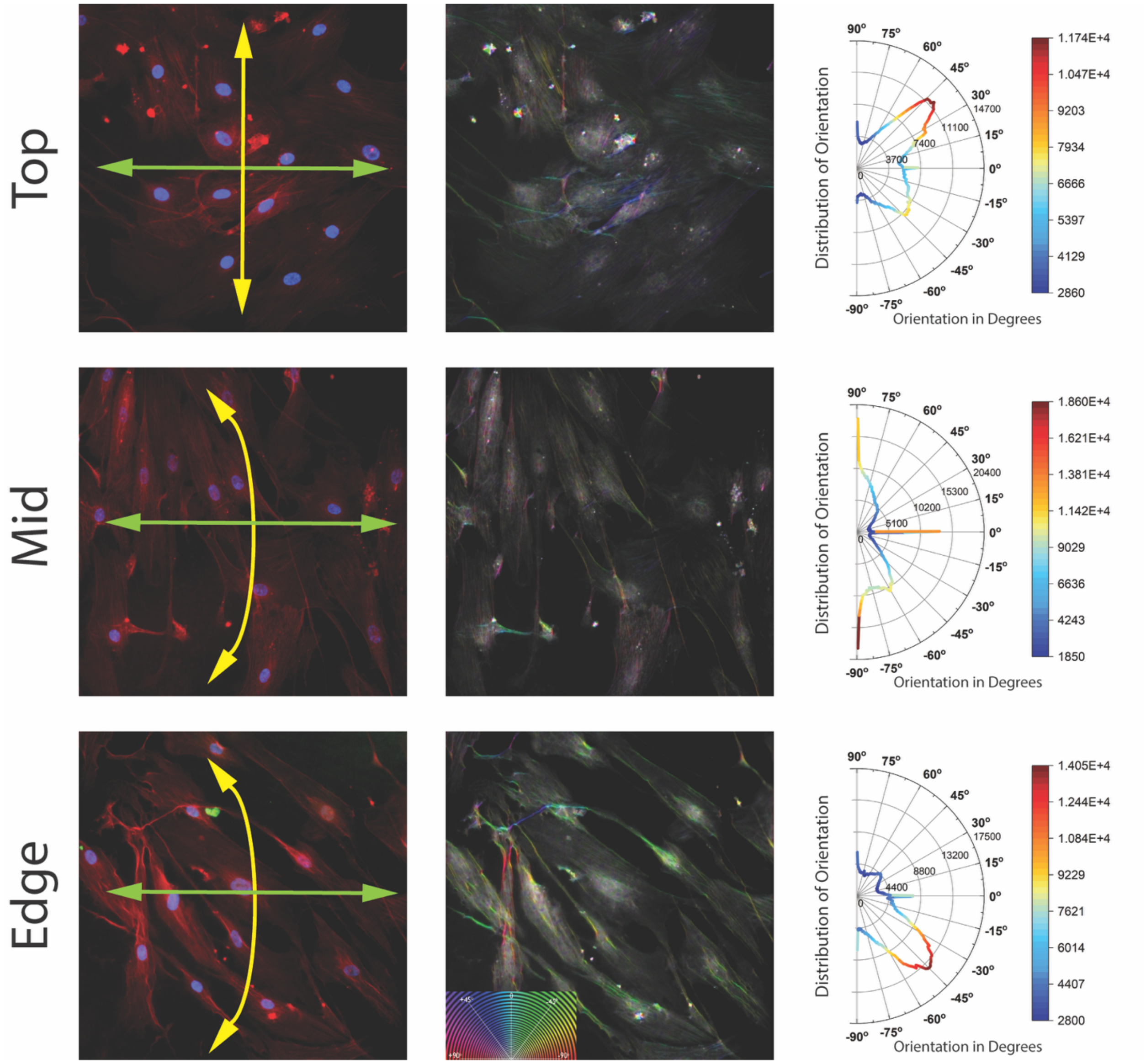


**Fig. S5.** Verification of trabecular meshwork cell orientation in different regions. Representative fluorescence images (left), orientation maps (middle), and corresponding orientation distribution plots (right) are shown for the top, mid, and edge regions. Yellow arrows showing the direction of hoop stress and green arrows showing the direction of meridional stress. Cells in the top region exhibit a largely random orientation distribution. In the mid region, the majority of cells are preferentially aligned along the hoop stress direction. In contrast, cells in the edge region predominantly align along the meridional stress direction, oriented toward the center, suggesting region-dependent cytoskeletal remodeling in response to local stress anisotropy.

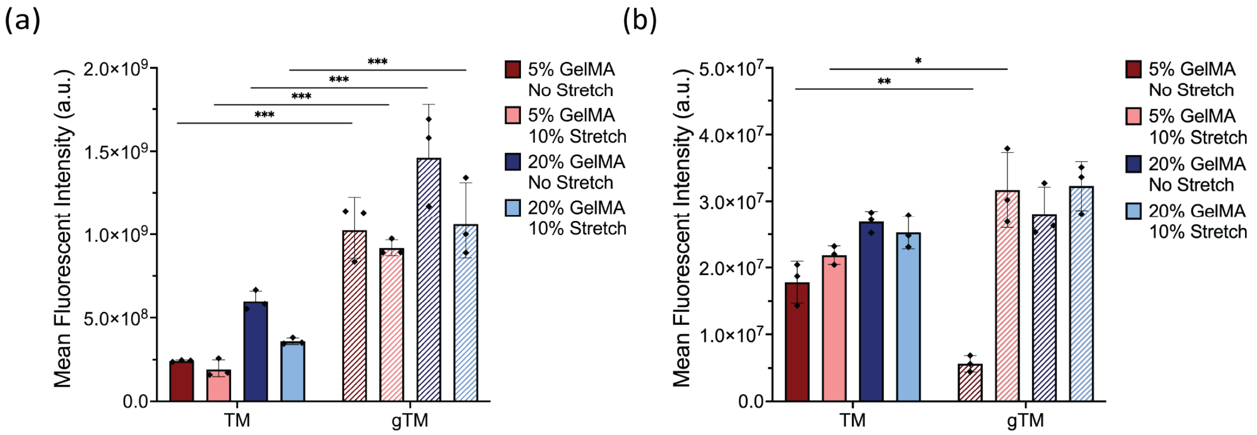


**Fig. S6. Comparative Analysis of Phenotypic Biomarker Expression between nTM and gTM Cells under varying matrix stiffness and mechanical Stretch.** Mean fluorescence intensity (a.u.) of α-SMA (a) and MYOC (b) comparing nTM and gTM cells within 5% and 20% GelMA hydrogels under no stretch or 10% stretch conditions. gTM cells exhibited significantly higher α-SMA expression compared to nTM cells across all conditions (all $p \leq 0.001$), whereas MYOC expression in gTM cells was significantly lower than nTM cells on compliant substrates under static conditions ($p \leq 0.01$) and significantly higher than nTM cells under combined 20% GelMA with 10% stretch ($p \leq 0.05$), reflecting stronger mechanical induction of MYOC in gTM cells. Data are presented as mean ± SD with individual biological replicates shown as black circles (n = 3). Statistical significance: *$p \leq 0.05$, **$p \leq 0.01$, ***$p \leq 0.001$.

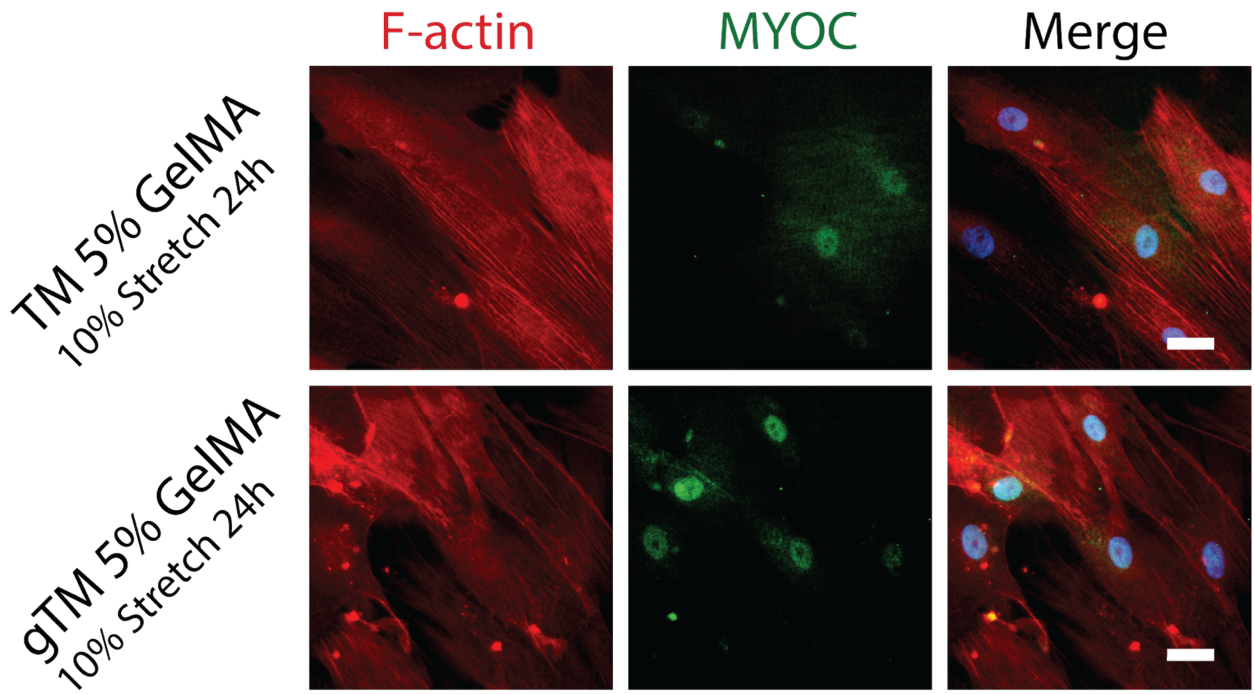


**Fig. S7. MYOC distribution in nTM and gTM cells under low stiffness and mechanical stretch.** Immunofluorescence comparison of MYOC localization in nTM and gTM cells cultured in 5% GelMA hydrogel under 10% stretch. F-actin (red), MYOC (green), and merged channels (with nuclei in blue) are shown for nTM cells (top row) and gTM cells (bottom row). In nTM cells, MYOC is distributed throughout both the cytoplasm and nucleus, whereas in gTM cells MYOC is concentrated within the nucleus, consistent with accumulation of misfolded MYOC. Scale bars: 50 µm.

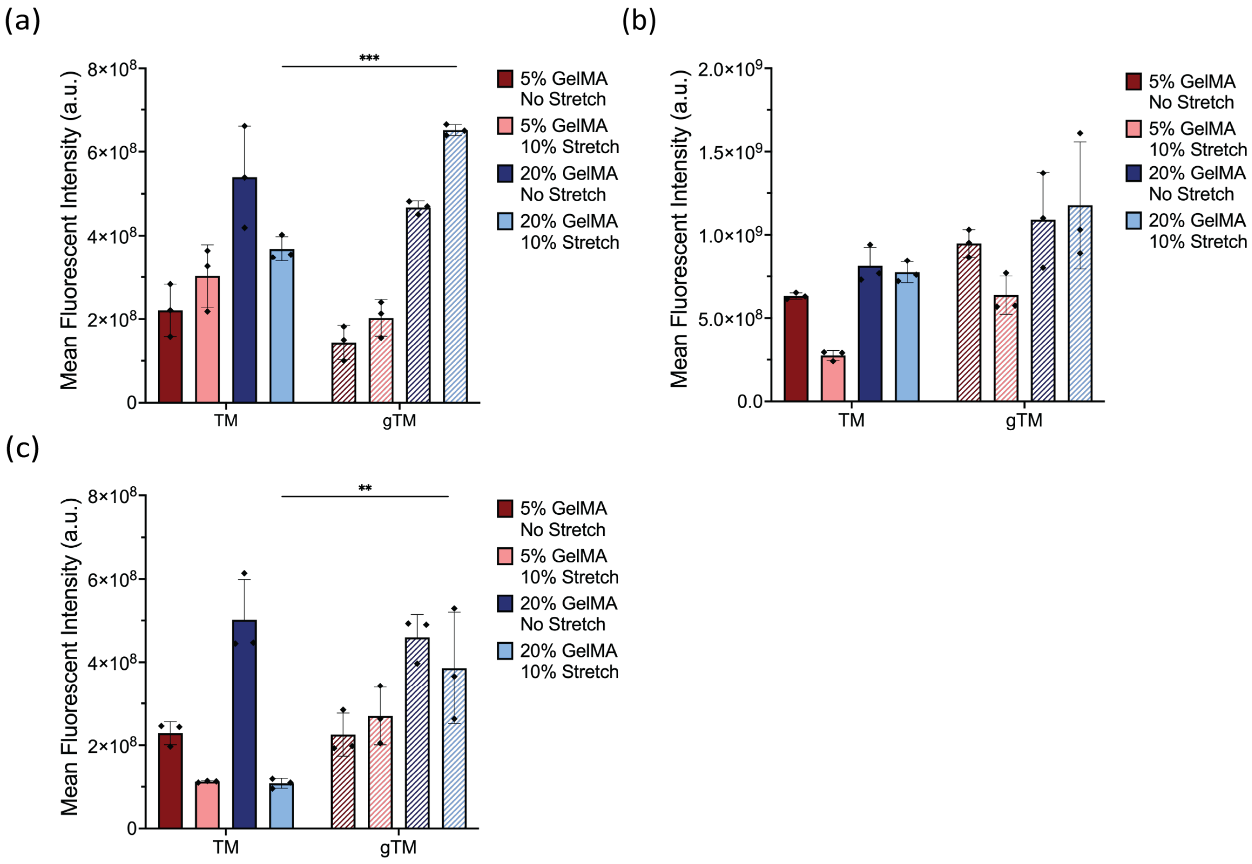


**Fig. S8. Comparative Analysis of Focal Adhesion and ECM Turnover Biomarker Expression between nTM and gTM cells under Varying Matrix Stiffness and Mechanical Stretch.** Mean fluorescence intensity (a.u.) of pFAK (a), MMP2 (b), and COL1 (c) comparing nTM and gTM cells within 5% and 20% GelMA hydrogels under no stretch or 10% stretch conditions. Red bars represent 5% GelMA (dark: no stretch; light: 10% stretch) and blue bars represent 20% GelMA (dark: no stretch; light: 10% stretch); solid bars indicate nTM cells and hatched bars indicate gTM cells. gTM cells exhibited significantly higher pFAK and COL1 expression compared to nTM cells specifically under the 20% GelMA with 10% stretch condition, suggesting that combined stiff and stretched conditions unmask cell-type differences in focal adhesion signaling and ECM remodeling. Data are presented as mean ± SD with individual biological replicates shown as black circles (n = 3). Statistical significance: $*p \leq 0.05$, $**p \leq 0.01$, $***p \leq 0.001$.

#### Equations

The arc length (L) was defined as:

$$L = \frac{\arctan\left(\frac{2h}{L_0}\right)(4h^2 + L_0)}{2h} \tag{1}$$

The change in arc length (ΔL) was calculated as:

$$\Delta L = L - L_0 \tag{2}$$

Linear strain (ε) was determined from:

$$\varepsilon = \frac{\Delta L}{L_0} \times 100\% \tag{3}$$

**Table S1.** Hyperelastic constitutive constants determined by iterative fitting of finite element simulations to experimental deflection data (PDMS density= $9.7 \times 10^{-7}$ kg/mm³ , bulk modulus= 20 MPa).

| Material | Material Model | Material Constant | Value |
|---|---|---|---|
| PDMS | Mooney-Rivlin | C1 (kPa) | 38.5 |
| | | C2 (kPa) | 2.93 |
| 5% GelMA | Veronda-Westmann | C1 (kPa) | 50 |
| | | C2 | 0.42 |
| 20% GelMA | Veronda-Westmann | C1 (kPa) | 60 |
| | | C2 | 0.46 |

### Three-way ANOVA Tests

#### Overall ANOVA of α-SMA

| | DF | Sum of Squares | Mean Square | F Value | P Value |
|---|---|---|---|---|---|
| Cell Type | 1 | 1.02993E19 | 1.02993E19 | 39.54552 | <0.0001 |
| Hydrogel Type | 1 | 6.03087E17 | 6.03087E17 | 2.31564 | 0.1376 |
| Stretch Type | 1 | 3.62828E17 | 3.62828E17 | 1.39313 | 0.24632 |
| Cell Type * Hydrogel Type | 1 | 1.1338E17 | 1.1338E17 | 0.43534 | 0.51396 |
| Cell Type * Stretch Type | 1 | 1.32314E14 | 1.32314E14 | 5.08039E-4 | 0.98215 |
| Hydrogel Type * Stretch Type | 1 | 2.77463E17 | 2.77463E17 | 1.06536 | 0.3095 |
| Cell Type * Hydrogel Type * Stretch Type | 1 | 1.53128E17 | 1.53128E17 | 0.58796 | 0.44866 |
| Model | 7 | 1.18033E19 | 1.68619E18 | 6.47438 | <0.0001 |
| Error | 33 | 8.59454E18 | 2.60441E17 | | |
| Corrected Total | 40 | 2.03979E19 | | | |

**At the 0.05 level, the population means of Cell Type are significantly different.**

**At the 0.05 level, the population means of Hydrogel Type are not significantly different.**

**At the 0.05 level, the population means of Stretch Type are not significantly different.**

**At the 0.05 level, the population means of Cell Type * Hydrogel Type are not significantly different.**

**At the 0.05 level, the population means of Cell Type * Stretch Type are not significantly different.**

**At the 0.05 level, the population means of Hydrogel Type * Stretch Type are not significantly different.**

**At the 0.05 level, the population means of Cell Type * Hydrogel Type * Stretch Type are not significantly different.**

#### Overall ANOVA of MYOC

| | DF | Sum of Squares | Mean Square | F Value | P Value |
|---|---|---|---|---|---|
| Cell Type | 1 | 1.28334E13 | 1.28334E13 | 1.23203 | 0.28342 |
| Hydrogel Type | 1 | 4.73926E14 | 4.73926E14 | 45.4976 | <0.0001 |
| Stretch Type | 1 | 3.98943E14 | 3.98943E14 | 38.29909 | <0.0001 |
| Cell Type * Hydrogel Type | 1 | 4.17384E13 | 4.17384E13 | 4.00695 | 0.06257 |
| Cell Type * Stretch Type | 1 | 2.88773E14 | 2.88773E14 | 27.72268 | <0.0001 |
| Hydrogel Type * Stretch Type | 1 | 2.84626E14 | 2.84626E14 | 27.32452 | <0.0001 |
| Cell Type * Hydrogel Type * Stretch Type | 1 | 9.78084E13 | 9.78084E13 | 9.38976 | 0.00741 |
| Model | 7 | 1.59865E15 | 2.28378E14 | 21.92466 | <0.0001 |
| Error | 16 | 1.66664E14 | 1.04165E13 | | |
| Corrected Total | 23 | 1.76531E15 | | | |

**At the 0.05 level, the population means of Cell Type are not significantly different.**

**At the 0.05 level, the population means of Hydrogel Type are significantly different.**

**At the 0.05 level, the population means of Stretch Type are significantly different.**

**At the 0.05 level, the population means of Cell Type * Hydrogel Type are not significantly different.**

**At the 0.05 level, the population means of Cell Type * Stretch Type are significantly different.**

**At the 0.05 level, the population means of Hydrogel Type * Stretch Type are significantly different.**

**At the 0.05 level, the population means of Cell Type * Hydrogel Type * Stretch Type are significantly different.**

#### Overall ANOVA of pFAK

| | DF | Sum of Squares | Mean Square | F Value | P Value |
|---|---|---|---|---|---|
| Cell Type | 1 | 1.08148E16 | 1.08148E16 | 0.17764 | 0.67623 |
| Hydrogel Type | 1 | 7.96684E17 | 7.96684E17 | 13.08618 | 0.00101 |
| Stretch Type | 1 | 3.25535E17 | 3.25535E17 | 5.34718 | 0.02734 |
| Cell Type * Hydrogel Type | 1 | 7.59124E15 | 7.59124E15 | 0.12469 | 0.72632 |
| Cell Type * Stretch Type | 1 | 2.44539E16 | 2.44539E16 | 0.40167 | 0.53073 |
| Hydrogel Type * Stretch Type | 1 | 1.43453E17 | 1.43453E17 | 2.35634 | 0.1346 |
| Cell Type * Hydrogel Type * Stretch Type | 1 | 4.03411E15 | 4.03411E15 | 0.06626 | 0.79851 |
| Model | 7 | 1.31257E18 | 1.8751E17 | 3.08 | 0.01347 |
| Error | 32 | 1.94815E18 | 6.08798E16 | | |
| Corrected Total | 39 | 3.26072E18 | | | |

**At the 0.05 level, the population means of Cell Type are not significantly different.**

**At the 0.05 level, the population means of Hydrogel Type are significantly different.**

**At the 0.05 level, the population means of Stretch Type are significantly different.**

**At the 0.05 level, the population means of Cell Type * Hydrogel Type are not significantly different.**

**At the 0.05 level, the population means of Cell Type * Stretch Type are not significantly different.**

**At the 0.05 level, the population means of Hydrogel Type * Stretch Type are not significantly different.**

**At the 0.05 level, the population means of Cell Type * Hydrogel Type * Stretch Type are not significantly different.**

#### Overall ANOVA of MMP2

| | DF | Sum of Squares | Mean Square | F Value | P Value |
|---|---|---|---|---|---|
| Cell Type | 1 | 5.60432E17 | 5.60432E17 | 8.24634 | 0.00719 |
| Hydrogel Type | 1 | 6.14603E17 | 6.14603E17 | 9.04343 | 0.0051 |
| Stretch Type | 1 | 2.70385E14 | 2.70385E14 | 0.00398 | 0.9501 |
| Cell Type * Hydrogel Type | 1 | 1.54773E16 | 1.54773E16 | 0.22774 | 0.63645 |
| Cell Type * Stretch Type | 1 | 3.32699E15 | 3.32699E15 | 0.04895 | 0.8263 |
| Hydrogel Type * Stretch Type | 1 | 7.7623E16 | 7.7623E16 | 1.14217 | 0.29319 |
| Cell Type * Hydrogel Type * Stretch Type | 1 | 8.7323E14 | 8.7323E14 | 0.01285 | 0.91046 |
| Model | 7 | 1.27261E18 | 1.81801E17 | 2.67506 | 0.02677 |
| Error | 32 | 2.17476E18 | 6.79613E16 | | |
| Corrected Total | 39 | 3.44737E18 | | | |

**At the 0.05 level, the population means of Cell Type are significantly different.**

**At the 0.05 level, the population means of Hydrogel Type are significantly different.**

**At the 0.05 level, the population means of Stretch Type are not significantly different.**

**At the 0.05 level, the population means of Cell Type * Hydrogel Type are not significantly different.**

**At the 0.05 level, the population means of Cell Type * Stretch Type are not significantly different.**

**At the 0.05 level, the population means of Hydrogel Type * Stretch Type are not significantly different.**

**At the 0.05 level, the population means of Cell Type * Hydrogel Type * Stretch Type are not significantly different.**

#### Overall ANOVA of Col1

| | DF | Sum of Squares | Mean Square | F Value | P Value |
|---|---|---|---|---|---|
| Cell Type | 1 | 5.18914E17 | 5.18914E17 | 18.85613 | 1.32662E-4 |
| Hydrogel Type | 1 | 2.68121E17 | 2.68121E17 | 9.74289 | 0.0038 |
| Stretch Type | 1 | 1.12369E17 | 1.12369E17 | 4.08323 | 0.05175 |
| Cell Type * Hydrogel Type | 1 | 3.27706E16 | 3.27706E16 | 1.19081 | 0.28332 |
| Cell Type * Stretch Type | 1 | 5.06995E16 | 5.06995E16 | 1.8423 | 0.18418 |
| Hydrogel Type * Stretch Type | 1 | 2.22817E17 | 2.22817E17 | 8.09666 | 0.00767 |
| Cell Type * Hydrogel Type * Stretch Type | 1 | 2.56845E16 | 2.56845E16 | 0.93332 | 0.34125 |
| Model | 7 | 1.23138E18 | 1.75911E17 | 6.39219 | <0.0001 |
| Error | 32 | 8.80629E17 | 2.75196E16 | | |
| Corrected Total | 39 | 2.112E18 | | | |

**At the 0.05 level, the population means of Cell Type are significantly different.**

**At the 0.05 level, the population means of Hydrogel Type are significantly different.**

**At the 0.05 level, the population means of Stretch Type are not significantly different.**

**At the 0.05 level, the population means of Cell Type * Hydrogel Type are not significantly different.**

**At the 0.05 level, the population means of Cell Type * Stretch Type are not significantly different.**

**At the 0.05 level, the population means of Hydrogel Type * Stretch Type are significantly different.**

**At the 0.05 level, the population means of Cell Type * Hydrogel Type * Stretch Type are not significantly different.**

**Table of Content**

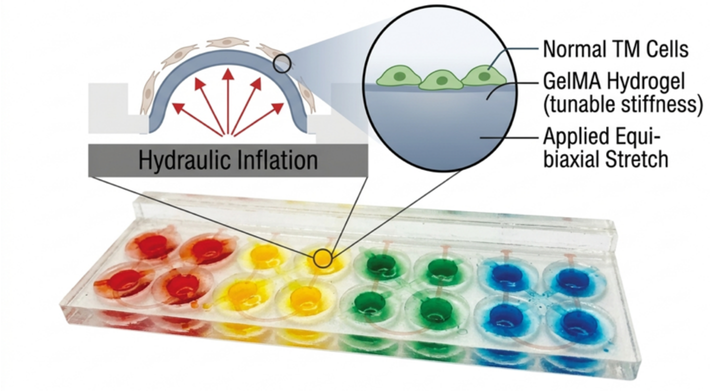


A hydrogel-integrated microfluidic platform simultaneously applies tunable substrate stiffness and equi-biaxial stretch to trabecular meshwork cells. Glaucomatous cells exhibit selective mechanotransduction dysregulation, including constitutively elevated contractile markers and impaired matrix remodeling, rather than generalized mechanosensory loss. Substrate stiffness and stretch interdependently regulate collagen expression, identifying tissue stiffening as a targetable driver of glaucoma-associated outflow dysfunction.
